# Deficits in Behavioral and Neuronal Pattern Separation in Temporal Lobe Epilepsy

**DOI:** 10.1101/2020.02.13.948364

**Authors:** A.D. Madar, J.A. Pfammatter, J. Bordenave, E.I. Plumley, S. Ravi, M. Cowie, E.P. Wallace, B.P. Hermann, R.K. Maganti, M.V. Jones

## Abstract

In temporal lobe epilepsy, the ability of the dentate gyrus to limit excitatory cortical input to the hippocampus breaks down, leading to seizures. The dentate gyrus is also thought to help discriminate between similar memories by performing pattern separation, but whether epilepsy leads to a breakdown in this neural computation, and thus to mnemonic discrimination impairments, remains unknown. Here we show that temporal lobe epilepsy is characterized by behavioral deficits in mnemonic discrimination tasks, in both humans (females and males) and mice (C57Bl6 males, systemic low-dose kainate model). Using a recently developed assay in brain slices of the same epileptic mice, we reveal a decreased ability of the dentate gyrus to perform certain forms of pattern separation. This is due to a subset of granule cells with abnormal bursting that can develop independently of early EEG abnormalities. Overall, our results linking physiology, computation and cognition in the same mice advance our understanding of episodic memory mechanisms and their dysfunction in epilepsy.

**Significance Statement:** People with temporal lobe epilepsy (TLE) often have learning and memory impairments, sometimes occurring earlier than the first seizure, but those symptoms and their biological underpinnings are poorly understood. We focused on the dentate gyrus, a brain region that is critical to avoid confusion between similar memories and is anatomically disorganized in TLE. We show that both humans and mice with TLE experience confusion between similar situations. This impairment coincides with a failure of the dentate gyrus to disambiguate similar input signals because of pathological bursting in a subset of neurons. Our work bridges seizure-oriented and memory-oriented views of the dentate gyrus function, suggests a mechanism for cognitive symptoms in TLE and supports a long-standing hypothesis of episodic memory theories.

## Introduction

Temporal lobe epilepsy (TLE) is characterized by recurring focal seizures originating in or near the hippocampus (Toyoda et al., 2013), microcircuit pathologies in brain regions including the hippocampus (Alexander et al., 2016) and memory-related cognitive deficits (Helmstaedter et al., 2003; Saling, 2009; Zhao et al., 2014). Although the hippocampus is a nexus for episodic memory that is critically affected during TLE, the relationship between TLE and hippocampus-dependent memory is insufficiently understood.

Episodic memory formation is thought to involve storage of neural representations in area CA3 of the hippocampus via Hebbian plasticity at recurrent excitatory synapses of coactivated cells (Rolls, 2010). In this view, a partial cue reactivates a subset of the CA3 ensemble that in turn recruits the other neurons of the original pattern resulting in recall of the original event. However, recurrent excitation is problematic because it theoretically a) predisposes the network to over-excitation that could trigger seizures (Traub and Wong, 1982; Le Duigou et al., 2014) and b) limits the number of patterns that can be stored without overlap (Rolls, 2010). Overlapping memory representations would in turn lead to interference during recall and thus cognitive confusion. To solve these problems, it was proposed that the dentate gyrus (DG) of the hippocampus acts as a) a gate and b) a pattern separator, so that similar cortical representations are transformed into sparse and dissimilar patterns before reaching CA3 (O’Reilly and McClelland, 1994; Hsu, 2007; Dengler and Coulter, 2016).

The function of DG as a gate for cortical activity to prevent seizure generation fails in TLE (Hsu, 2007; Krook-Magnuson et al., 2015; Dengler and Coulter, 2016; Lu et al., 2016). Granule cells (GCs), the output neurons of DG, lose their usual sparseness due to network reorganization (Artinian et al., 2011; Dengler and Coulter, 2016; Dengler et al., 2017) making cortical excitation easier to propagate to CA3 (Behr et al., 1998; Patrylo et al., 1999; Ouedraogo et al., 2016). Even in healthy animals, repeated excitation of GCs induces seizures (Krook-Magnuson et al., 2015) and prolonged stimulation of DG causes TLE (Sloviter, 1983).

An alternate view of the DG function, developed in parallel to the concept of dentate gate, is that it performs pattern separation: the transformation of similar cortical patterns into dissimilar hippocampal representations. This process theoretically supports mnemonic discrimination, the ability to distinguish between similar memories. Indeed, DG lesions impair mnemonic discrimination both in rodents (Treves et al., 2008; Kesner and Rolls, 2015; Kesner et al., 2016) and humans (Yassa et al., 2011; Baker et al., 2016; Bennett and Stark, 2016; Dillon et al., 2017). Moreover, computational models (Chavlis and Poirazi, 2017) and recent experiments (Berron et al., 2016; Knierim and Neunuebel, 2016; Madar et al., 2019a, 2019b; Woods et al., 2020) suggest that the DG network supports multiple forms of pattern separation (Santoro, 2013). Whether such computations underlie mnemonic discrimination remains unclear.

Unsurprisingly, TLE negatively impacts hippocampal-dependent memory in humans (Coras et al., 2014) and rodents (Gröticke et al., 2008; Chauvière et al., 2009; Müller et al., 2009; Inostroza et al., 2013; Lenck-Santini and Scott, 2015). However, the effect of TLE specifically on DG computations and DG-dependent cognition is understudied. It was only recently reported that patients with TLE are impaired on mnemonic discrimination tasks (Reyes et al., 2018; Poch et al., 2020; Lalani et al., 2021) and that TLE causes deficits in DG-dependent object location memory in mice (Bui et al., 2018; Kahn et al., 2019). A computational model has also suggested that hippocampal pathologies in TLE would degrade DG pattern separation (Yim et al., 2015) but the hypotheses that 1) TLE causes a breakdown in DG neural pattern separation and 2) that such a failure causes mnemonic discrimination impairments remain experimentally untested.

Here we investigated, in humans and mice, whether TLE is characterized by deficits in mnemonic discrimination and then recorded, in brain slices from the same mice, the spiking patterns of single GCs in response to parametrically varied afferent stimulation in order to gauge neuronal pattern separation.

## Materials and Methods

### Human Behavior

A mnemonic similarity task ***(Stark et al., 2019)***, also known as a behavioral pattern separation (BPS) task ***(Stark et al., 2013)***, was administered to 15 patients in the University of Wisconsin-Madison Epilepsy Monitoring Unit. Under an approved Institutional Review Board protocol and after obtaining informed consent, the task was administered in conjunction with a standard neuropsychiatric evaluation and during electroencephalographic (EEG) recording, both of which are part of standard practice for diagnosing the patients’ seizures. Patients were 18-65 years old, male and female. Only patients with a preliminary diagnosis of TLE were included in our analysis. As controls, 20 subjects without epilepsy, recruited to match the patients’ age and sex distributions (family members of the patients when possible, or other volunteers), were also tested. Consent documents, medical records and primary data are on file in a secure location within the Dept. of Neurology. Data were deidentified prior to analysis. Subjects received no compensation for participation.

The visual, object recognition-based BPS task is described in Yassa et al. (2011) and has been further validated by demonstrating mnemonic discrimination deficits during normal aging and multiple neurologic and psychiatric disorders (Stark et al., 2019) recently including TLE (Lalani et al., 2021). It was implemented on a laptop by trained neuropsychiatric postdoctoral fellows (J.B. and E.I.P.) using software distributed freely by the Stark lab (http://faculty.sites.uci.edu/starklab/mnemonic-similarity-task-mst/). Briefly, participants viewed a series of pictures of everyday objects: During Phase 1, 128 different images were presented and the subject was asked to classify each as “indoor” or “outdoor”, simply to engage the subject’s attention. During Phase 2, a new series of 192 images was presented: 1/3 repeated from Phase 1 (*Repeated*), 1/3 new but similar to images from Phase 1 (*Lure*), and 1/3 new and completely different from Phase 1 (*Novel*). Participants were asked to classify each image of the Phase 2 set as “Old”, “Similar”, or “New”. BPS was evaluated with a *discrimination score* computed as p(“Similar”|Lure) - p(“Similar”|Novel), as in past research (Yassa et al., 2011; Stark et al., 2019). We also computed a *recognition score* as p(“Old”|Repeated) - p(“Old”|Novel), which measures performance for an “easy” mnemonic discrimination not necessarily dependent on the hippocampus (Kirwan et al., 2012; Lalani et al., 2021).

### Mouse Experiments

Male mice (C57BL6J, 5-6 weeks old) were received from Envigo (formerly Harlan, Madison, WI), housed in groups and allowed to acclimate to their new home environment for one or two weeks. Animals then underwent epilepsy induction with kainic acid (KA) injections (see below) (J.A.P.). A control group of mice was injected with saline and another received no injection: both were pooled together and considered as the control group because our analyses did not reveal any difference. 7-9 weeks after injection, mice started 4 weeks of behavioral testing on the BPS task. After completion of behavioral testing, animals (~18-20 weeks old) were transferred to a different building for EEG implantation. Each animal was recorded continuously for three days before being transferred back to the original building and sacrificed for slice electrophysiology (~20-24 weeks old). After each building-to-building transfer, mice were allowed a period of acclimation ranging from two days to two weeks. Behavioral testing (J.A.P, S.R. and M.C.), EEG recordings (E.P.W.) and slice electrophysiology (A.D.M) were performed by different researchers, all of whom were blind to KA or Control treatments.

### Epilepsy induction

During the induction process, when not being handled, mice were individually housed in enclosed ~150 cm^3^ acrylic cubicles with opaque sides and clear front portals with holes to allow air exchange, food and bedding. Animals were then randomly assigned to KA or Control treatment, ear punched for identification and weighed. Animals were induced for epileptogenesis using the repeated low-dose kainate method *(Hellier et al., 1998; Sharma et al., 2018)*: a series of intraperitoneal injections based on the following schedule. Animals first received 10 mg/kg of kainic acid (Tocris Bioscience, UK) mixed in 1x PBS (8 ml per 10 mg) prepared from PBS tablets (Dot Scientific, Michigan) with deionized distilled water and filter sterilized (Millipore, Burlington, MA). Control animals were injected with an equivalent volume of saline made of 1x PBS. Animals continued to receive injections at 5mg/kg every 20 minutes until status epilepticus (SE) occurred (same number of saline injections in control mice). Some animals received alternating 5mg/kg and 2.5 mg/kg injections after the initial 10 mg/kg dose, but we found this schedule took more time than 5 mg/kg injections and our survival rate (>90% across all cohorts) was no different between the two schedules. Animals were considered to be in SE when displaying persistent behavioral seizures of level 4-5 on the Racine Scale (Racine, 1972), less than 5 minutes apart, for a minimum of 30 minutes. Animals received 4-9 injections depending on tolerance to kainic acid and injection schedule. After the injection schedule, animals were given fresh apple slices, monitored until SE ceased (assessed from normal posturing, within 1-1.5 hours), and returned to group housing. They were monitored, weighed and animals weighing less than their preinjection levels were given 0.4 ml of 1x PBS via intraperitoneal injection on a daily basis until their weight exceeded preinjection level. No animal injected with saline ever experienced status epilepticus, lost weight, or required recovery injections of 1x PBS. Mice remained in the vivarium for 7-9 weeks after injections, allowing time for the development of epilepsy in KA mice - the latent period in the KA model of TLE has previously been evaluated as 10-30 days before the first spontaneous electrographic seizure, and ~11±5.4 weeks before the first spontaneous convulsive seizure *(Lévesque and Avoli, 2013)*. During this time two mice (1 KA and 1 control) became aggressive and were removed from group housing to be caged individually, but no effect of housing on our results was detected.

### Mouse Behavior

At 14-16 weeks of age, mice started behavioral testing on a variant of a novelty recognition-based BPS task that is well established in rodents to report mnemonic discrimination abilities *(van Hagen et al., 2015)*. Our protocol lasted 4 weeks, including 1 week of habituation and 3 weeks of trials. During the first week of habituation, each mouse was gently handled by the experimenter for five minutes every day. On the second week mice were split into three groups (A, B, and C). Each group went through a three-day schedule where animals were habituated on the first and second days and performed the object location task on the third day. Groups were staggered by one day each, such that group A was scheduled Monday-Wednesday, B Tuesday-Thursday and C Wednesday-Friday. Animals were housed following a 12:12 light-dark cycle and always handled or tested between 1 pm and 4pm (light period).

The arena used for the BPS task was a 63.5 x 63.5 x 15 cm open topped Plexiglas box wrapped on the outside and bottom with black felt cloth. On each side of the arena were different images serving as proximal cues for orientation. The room also had numerous distinct distal cues (e.g. the video camera placed over the center of the arena, shelving, etc.), that were kept unchanged. During behavioral testing, the room was lit with a single bank of overhead fluorescent lamps.

For the BPS task, animals underwent three phases of exploration in the arena, each lasting 3 min and separated by 1.5 min of single housing in a solitary chamber. In Phase 1 (habituation) the arena was empty, in Phase 2 (sampling) the arena contained two identical objects in the center, and in Phase 3 (testing) one object had been randomly selected and shifted by a distance of 7, 14 or 21 cm. The arena and objects were cleaned with 70% ethanol before and after each phase. During Phases 2 and 3, the experimenter recorded the amount of time that the mouse spent interacting with each object with two silent stopwatches. The mouse was considered to be interacting with an object if it was within 1 cm of it and oriented towards, sniffing, scratching, or on top of the object. These three phases were repeated for a different object shift each week (7, 14 or 21 cm), with the order of shifts randomized. Objects used for the task were ~6 cm tall plastic colored figures or metal cylinders, with a footprint of ~2 cm in diameter.

For a subsample of experiments, trials were recorded with a 1080p webcam positioned above the arena. During all phases, the experimenter sat in the corner of the room out of the line of sight of the mouse. The experimenter was consistent in appearance and smell (e.g., same experimenter for a given cohort, wearing same white lab coat and gloves of the same color each day; use of scented soap, etc. was minimized).

### BPS Analysis

The discrimination ratio for each trial was computed as (T_moved_ – T_unmoved_) / T_total_ where T_moved_ is the time spent exploring the moved object, T_unmoved_ is the time spent exploring the unmoved object and T_total_ is the sum of T_moved_ and T_unmoved_. To visualize individual discrimination performance over all distances, each animal tested for all three object shift conditions was considered as a point with coordinates corresponding to that animal’s discrimination for each object shift. An individual Discrimination Impairment score was computed as the Mahalanobis distance from the centroid of the group of control mice *(De Maesschalck et al., 2000)*: 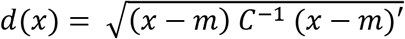, where x is a vector of coordinates corresponding to an animal, m is the vector of coordinates of the control centroid and C is the covariance of the control distribution (The impairment score thus represents the distance of point x from the control centroid in number of standard deviation of the control distribution). The exploration times used to calculate the reported discrimination ratios and impairment scores were manually collected by experimenters during the BPS task, and experiments with exploration times < 4 s on either phase 2 or 3 were excluded of all analyses *(Pofahl et al., 2021)* except for computing the impairment score over all object shifts. We (M.V.J. and J.A.P.) also developed a home-written program using video recordings for motion-tracking and trajectory analysis of each mouse during behavioral testing. Discrimination ratios computed from automated tracking data were well correlated to the ratios from manually recorded times, and led to similar results (R^2^ = 66%, *F(1, 93)* = 16.07, *P* < 0.0001). Automated tracking data were used to assess a) the ambulatory activity (distance travelled / time travelled) and b) thigmotaxis (the tendency to stay close to walls) as an estimate of anxiety. Thigmotaxis *(Simon et al., 1994)* was evaluated as the proportion of time spent within 6.35 cm of the wall (10% of the arena width).

### Mouse electroencephalography (EEG)

EEG electrode implantation was performed at ~18-20 weeks of age for all animals, following a previously established protocol *(Wallace et al., 2015)*. Briefly, mice were anesthetized with isoflurane and stainless steel screw electrodes were implanted in the skull (bregma +1.5 mm and 1 mm right, bregma −3 mm and 1 mm left, and lamda −1 mm at midline). Two stainless steel braided wires were placed in the nuchal muscles for electromyography (EMG) recording. After a 72 hour recovery, we transferred mice into individual tethered EEG acquisition chambers and allowed a >12 hour acclimation period. We acquired EEG and EMG signals continuously for 3 days. Recordings were digitized with an XLTek amplifier (XLTEK, USA) sampled at 1024 Hz. Ad libitum access to food and water was ensured. Note that the resulting EEG dataset was previously reported and analyzed in Pfammatter et al. (2018).

### Interictal spike (IIS) analysis

In humans and most animal models of acquired TLE, overt seizures are relatively rare (often much less than once per day) *(Lévesque et al., 2016)*. In contrast, nonconvulsive and subclinical epileptiform events such as interictal spikes (IISs) can be very frequent, up to hundreds per day. Therefore, in order to assess epileptiform activity in KA animals, we used a modification of our previously published principal components (PC)-based method to quantify IISs *(Pfammatter et al., 2018)* and calculated an “Hourly IIS index” for each animal. All detected high-amplitude EEG events from 9 Ctrl and 15 KA animals were projected into the space spanned by their first three PCs. The main modification here is that, instead of using a Gaussian Mixture Model (GMM) to assign events to ‘clusters’, we simply gridded the PC space into ‘voxels’ and computed relevant quantities within each voxel exactly as we computed those same quantities within GMM ‘clusters’ previously: 1) The probability that events within a voxel are characteristic of an epileptogenic treatment (i.e., KA) was computed as the voxel-wise proportion of events coming from KA mice, 2) Only the voxels above chance level were considered specific to epileptogenic treatment and probabilities were scaled accordingly, 3) The Hourly IIS Index of a given animal was defined as the average frequency of detected events weighted by the scaled probabilities of each event’s voxel. This new voxel-based method has the advantages to make no assumptions about the structure of the data in PC space or about the number of clusters to fit. Instead, the main free parameter is now the voxel volume: we selected a size of 10 cubic PC units following the same optimization procedure described in Pfammatter et al. (2018) in order to avoid overfitting. Extensive exploration (not shown) revealed that the final results are similar to the GMM method and are not importantly affected by moderate changes in voxel volume.

### Slice electrophysiology

Mice used for electrophysiology were p115-p182 at the time of experiment (mean ± SEM: p141 +/- 4 days; no difference in age between treatments: KA n = 13, Ctrl n = 6, U-test: P = 0.9, Z = 0.2, rank sum = 246). Age was also not correlated with any summary statistics presented in this study.

Adult mice were euthanized by transcardial perfusion with oxygenated PBS under isoflurane anesthesia, before decapitation and brain extraction. 400 μm horizontal slices of the ventral and intermediate hippocampus were prepared as detailed in Madar et al. (2019a). After slicing in a sucrose-based cutting solution *(Yi et al., 2015)*, slices were transferred to an incubation chamber filled with 50% cutting solution and 50% artificial cerebrospinal fluid (aCSF) at 37°C for 30 minutes, then room temperature. Patch-clamp recordings were done in a chamber submerged with aCSF containing (in mM) 125 NaCl, 25 NaHCO3, 2.5 KCl, 1.25 NaH2PO4, 2 CaCl2, 1 MgCl2, and 25 D-Glucose, flowing at 5 ml/min and saturated with a gas mixture of 95% O2 and 5% CO_2_. Stimulation was applied through a double-barreled “theta” pipette filled with aCSF. Patch pipettes were filled with an intracellular solution of the following composition (in mM): 135 K-gluconate, 5 KCl, 0.1 EGTA, 10 HEPES, 20 Na-Phosphocreatine, 2 Mg_2_-ATP, 0.3 Na-GTP, 0.25 CaCl_2_ adjusted to pH 7.3 with KOH and 310 mOsm with H_2_O, leading to a 2-5 MΩ pipette resistance in aCSF. Whole-cell patch-clamp recordings of single DG GCs in response to electric stimulation of the outer molecular layer (perforant path) were performed as detailed in Madar et al. (2019a), and the stimulation protocols used to test neural pattern separation were the same as in Madar et al. (2019b). Briefly, input sets used for stimulation were composed of five (type 1) or ten (type 2 and 3) spiketrains (two seconds long), delivered sequentially (separated by five seconds of pause) and repeated ten or five times, respectively, in order to yield fifty output spiketrains. The stimulation pipette was placed >100μm lateral to the recorded GC to avoid direct stimulation of GC dendrites, with the baseline membrane potential held at −70 mV for current and voltage-clamp recordings.

Intrinsic electrophysiological properties of recorded GCs were the following (mean ± SEM for Ctrl / KA): resting membrane potential V_rest_ = −78.0 ± 1.9 / −81.4 ± 1.2 mV; membrane resistance R_m_ = 139 ± 17 / 178 ± 16 MΩ; membrane capacitance C_m_ = 18 ± 1.2 / 17 ± 0.7 pF. There were no significant differences between control and KA mice (U-tests: P = 0.1, 0.2, 0.7; Z = 1.6, - 1.2, 0.4; rank sums = 294, 198.5, 252.5 respectively).

### Neural pattern separation analysis

Pattern separation levels were calculated as in Madar et al. (2019a, b). The general approach is to divide each spiketrain into time bins of a specific duration τw, then consider a certain set of features of this vectorized spiketrain as the relevant neural code (e.g. the number of spikes in a bin), and measure the similarity between pairs of spiketrains: the amount of pattern separation (or convergence, for negative values) is given by the difference between the similarity of two input spiketrains and the similarity of the corresponding pair of output spiketrains. We used six different metrics that considered distinct neural codes (i.e. each metric focuses on different spiketrain features): two similarity metrics sensitive to binwise synchrony between pairs of spiketrains (the Pearson’s correlation coefficient R and the normalized dot product NDP), a similarity metric sensitive to variations in firing rate and burstiness (the scaling factor SF), two dispersion metrics sensitive to complementary aspects of burstiness (Compactness and Occupancy) and a dispersion metric simply considering the overall firing rate of a spiketrain (FR) (see **Table 1**). NDP and SF are two complementary metrics that consider spiketrains as vectors of spike count per bin: NDP is the cosine of the angle between vectors (i.e. measuring the degree of orthogonality) whereas SF is the ratio of norms (i.e. measuring length differences). R considers vectors of [spike count – mean spike count], which loses the original angle between the vectors of raw spike counts and leads to considering common periods of silence as correlated, unlike NDP. Compactness is given by: 1 - proportion of time bins with at least one spike. Occupancy is the average number of spikes in bins with at least one spike. FR is simply the average firing rate over the duration of the spiketrain. See Madar et al. 2019b for a detailed discussion of each metric.

**Table 1.** Neural code assumed by each similarity/dispersion metrics.

|  | R | NDP | SF | Burstiness 1:<br>Compactness | Burstiness 2:<br>Occupancy | FR |
| --- | --- | --- | --- | --- | --- | --- |
| Spike count code | X | X | X |  | X | X |
| Excludes common periods of silence |  | X |  | X | X |  |
| Sensitive to number of spikes in corresponding time bins (binwise synchrony) | X | X |  |  |  |  |
| Sensitive to firing rate or burstiness |  |  | X | X | X | X |
| Sensitive to number of occupied bins |  |  | X | X |  |  |
| Sensitive to average number of spikes per bin |  |  | X |  | X |  |
| Sensitive to total number of spikes (i.e. spike count in a single bin of 2s) |  |  | X |  |  | X |

Compactness, Occupancy and FR are not similarity metrics, they simply quantify a given spiketrain feature F for each spiketrain. Instead of measuring the similarity between spiketrains we thus quantified the absolute difference between two spiketrains in terms of F (e.g. |FR of spiketrain A – FR of spiketrain B|). For a set of multiple spiketrains, we computed the dispersion D as the mean value of pairwise absolute differences of F over all spiketrains (excluding self-comparisons and, in the case of the output sets, all comparisons between output spiketrains resulting from the same input spiketrain). The degree of pattern separation was thus D_output_ - D_input_.

When using a similarity metric S (R, NDP or SF), pattern separation was assessed following the same logic. First, the similarity between each pair of spiketrains of a given set was computed: S_input_ was the average for all pairs of an input set (excluding self-comparisons), and S_output_ was the average of all pairs in an output set (excluding comparisons between output spiketrains coming from repetitions of the same input train). The degree of pattern separation was thus S_input_ - S_output_. To gain a finer view, we also performed a pairwise analysis (as in Madar et al. 2019b) where each of the pairwise S_input_ of an input set were distinguished (10 pairs for input sets of type 1, 45 pairs for input sets of type 2 and 3). The pairwise S_output_ (pw S_output_) was thus the average similarity across all pairs of output spiketrains resulting from a given pair of input spiketrains. Pattern separation was computed as pw S_input_ – pw S_output_.

### Experimental design and statistical analysis

Sample sizes correspond to the standards of each field (human behavior, mouse behavior, slice electrophysiology) such that large effects could be detected. Data analysis was performed using MATLAB (Mathworks, Natick, MA, USA). The Lilliefors test was used to verify the normality of data distributions ***(Lilliefors, 1967)***. Parametric or nonparametric statistical tests were appropriately used to assess significance (p-value < 0.05). Throughout the Results section, KW ANOVA corresponds to the nonparametric Kruskal-Wallis analysis of variance, U-test corresponds to the Wilcoxon rank-sum test, and KS test corresponds to the two-sample Kolmogorov-Smirnov test (sidedness is specified in the legends).

To analyze performance of mice on the BPS task we performed a two-way ANOVA with the Matlab function *anova* based on a linear mixed-effects model built using *fitlme*, with the distance of the moved object as a continuous fixed effect, animal treatment (Ctrl vs. KA) as a categorical fixed effect, and animal identity nested within object distances as a random effect to account for repeated measurements. The same procedure was used to test for differences in object exploration times, ambulatory activity and thigmotaxis. For bootstrapped estimates and confidence intervals, we used the Matlab function bootci with 1000 bootstrap samples and Bca method.

In the pattern separation analysis, to determine whether (S_input_, S_output_) distributions were significantly different, we performed an analysis of the covariance (ANCOVA) using separate parabolic or linear regression models, implemented in MATLAB with a custom-written code following the method described in (Motulsky and Ransnas, 1987), as in Madar et al. (2019a, b)

For the non-linear regression between pBurst and pattern separation, we used the Matlab function *fitnlm* with a three-parameter model function *f*(*x*) = *a*/(*x* + *b*) + *c*.

Violin plots were generated using code from Bastian Bechtold (2016): https://github.com/bastibe/Violinplot-Matlab, DOI: 10.5281/zenodo.4559847

### Code accessibility

Raw data for all figures are available at: https://www.ebi.ac.uk/biostudies/studies/SBSST353. Code to read and analyze the data is available at: https://github.com/jessePfammatter/DeficitsBehavNeuronalPatternSep and https://github.com/antoinemadar/PatSepSpikeTrains

## Results

### Behavioral pattern separation deficits in TLE

Mnemonic discrimination is often called behavioral pattern separation (BPS) because it is hypothesized to be the behavioral outcome of neural pattern separation. It is the ability to discriminate between similar experiences (event, environment, object, etc.) that occurred at different times ***(Santoro, 2013)***.

We first investigated the effect of TLE on mnemonic discrimination in humans. We used the *Mnemonic Similarity Task* developed by the Stark lab, an established DG-dependent BPS task (Baker et al., 2016; Stark et al., 2019) where participants must distinguish between similar images presented at different times (**Figure 1A, Materials and Methods – Human behavior**). Patients previously diagnosed with TLE showed a severe deficit (~50%) compared to nonepileptic subjects in their ability to correctly identify objects as being similar but not identical (**Figure 1**). We also measured basic recognition memory, which relies on multiple temporal lobe areas and is not necessarily dependent on the hippocampus (Eichenbaum et al., 2007; Kirwan et al., 2012): patients with TLE had trouble recognizing old images and discriminating them from new and completely different images, but recognition deficits were not predictive of the deficits on the more challenging DG-dependent discrimination task (**Figure 1E**). This is consistent with the emerging view of TLE as a disorder with distributed brain abnormalities causing a variety of distinct cognitive impairments (Bell et al., 2011). These results support our hypothesis that TLE impairs DG-dependent mnemonic discrimination, thus warranting further study of the impact of TLE on DG computations that might explain this cognitive deficit.

**Figure 1.**
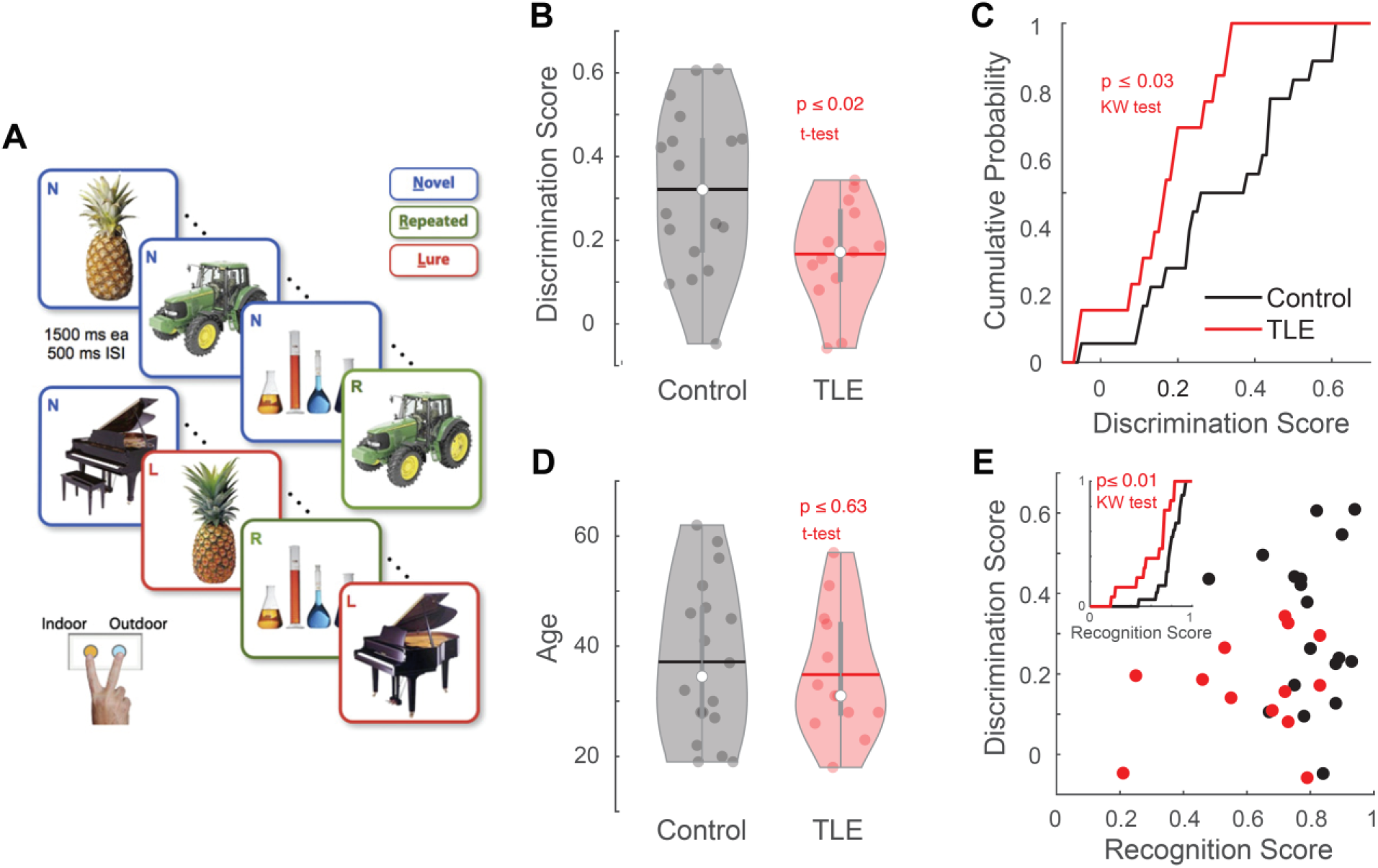
Human patients with TLE have mnemonic discrimination deficits. **(A)** A schematic (adapted with permission from Yassa et al., 2011) of the mnemonic discrimination task given to patients with TLE and nonepileptic control subjects. The discrimination score measures the ability of participants to correctly identify images as similar but not identical to a previously seen image. **(B)** Violin plots (open circle = median, horizontal bar = mean, scattered dots = subjects, grey vertical bar = quartiles) show the distribution of individual mnemonic discrimination scores, computed as p(“Similar”|Lure) - p(“Similar”|Novel). Patients with TLE (red, n = 13) display a deficit compared to nonepileptic control subjects (black, n = 18). Unpaired two-sided T-test: P = 0.017, T(29) = 2.54. **(C)** The same data from B, presented as cumulative frequency distributions. A non-parametric one-way KW ANOVA on the indices grouped by treatment confirms a highly significant deficit in visual pattern separation memory for human subjects with TLE (P = 0.031, χ^2^(1,29) = 4.67). **(D)** TLE and Ctrl groups were properly age matched (unpaired two-sided T-test: P = 0.63, T(29) = 0.48). There were slightly more women than men in the control group, but the gender difference was not significant between the two treatments (one-way KW ANOVA on gender values, 1 or 2, grouped by treatment: P = 0.052, χ^2^(1,29) = 3.77). **(E)** Inset: Cumulative frequency distributions of recognition scores, computed as p(“Old”|Repeated) - p(“Old”|Novel). Patients with TLE (red) have a lower recognition score than control subjects (black). One-way KW ANOVA on the indices grouped by treatment: P = 0.006, χ^2^(1,29) = 7.64. Main panel: For patients or control subjects, there is no apparent relationship between recognition and discrimination scores. Note that some patients with TLE can have a normal recognition score but a discrimination score close to 0.

To investigate the effect of TLE at the behavioral, computational and cellular levels, we turned to the common kainate animal model of acquired TLE (KA, **Methods – Mouse experiments**). An automated detection algorithm of interictal spikes (IISs) allowed us to quantify epileptiform activity in each animal even in the absence of recorded seizures (**Methods – IIS analysis** and see Pfammatter et al. 2018). Although we observed seizures in only two KA animals (Pfammatter et al. 2018), most KA mice developed IISs (**Figure 2**), demonstrating epileptogenesis. The detected IISs belonged to two distinct groups (**Figure 2D**), both having waveform features typical of epilepsy (**Figure 2E-F**) (White et al., 2010; Chauvière et al., 2012; Huneau et al., 2013; Pfammatter et al., 2018).

**Figure 2.**
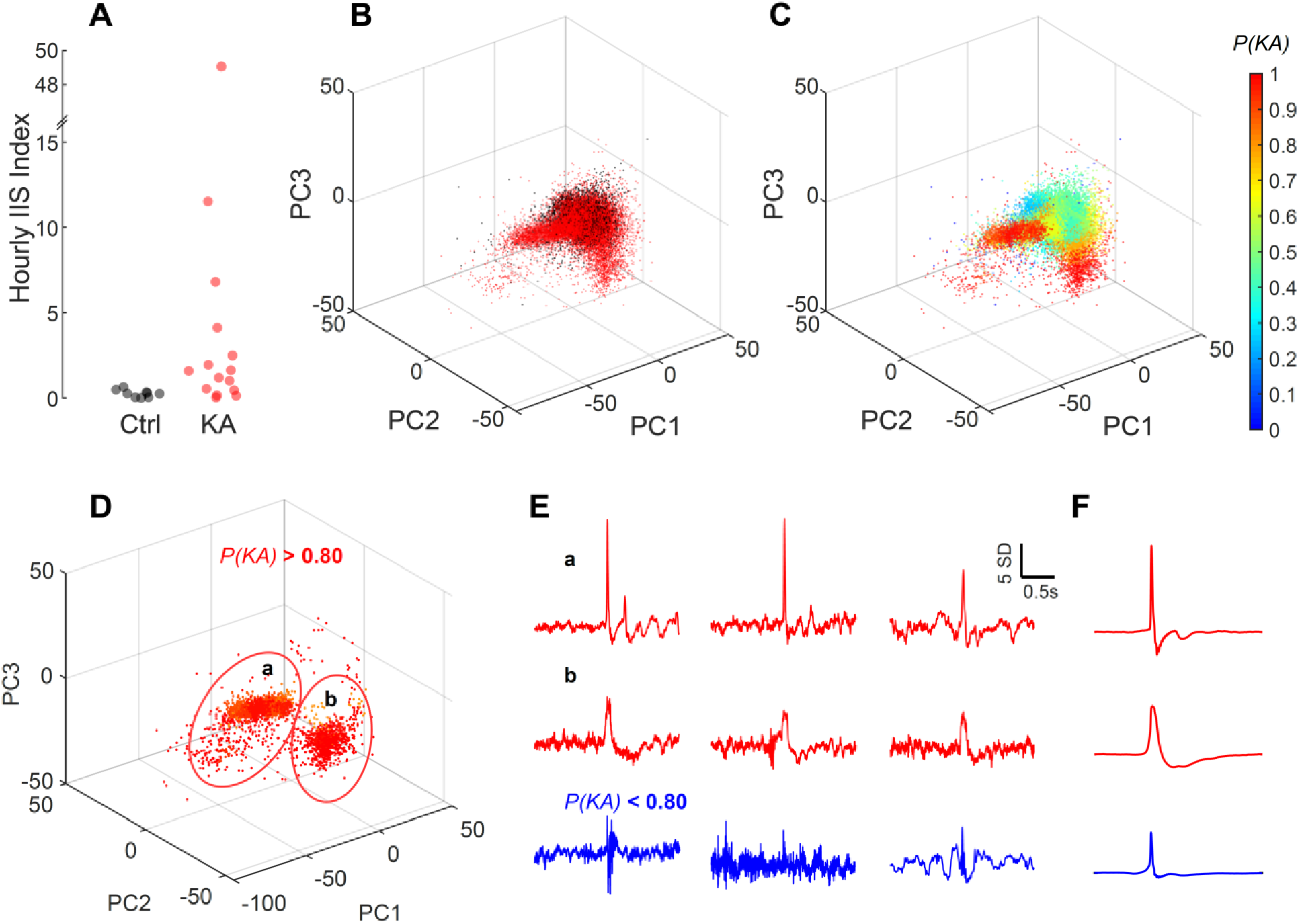
Kainate-injected mice have epileptiform electrographic events with a range of event frequency. **(A)** The Hourly Interictal Spikes (IIS) index, a proxy for epilepsy severity based on EEG recordings (Pfammatter et al., 2018), was calculated for 9 Ctrl (black) and 15 KA (red) animals. KA animals have a significantly higher Hourly IIS index than Ctrl animals (U-test: *P* = 0.009, Z = 2.207, rank sum = 225). **(B)** To compute the index, high-amplitude events (200 ms around a spike) were identified from left frontal EEG and projected on the first three principal components (PC) of the ensemble of events from all mice. Each dot here is an EEG event, red when from a KA animal and black when from a Ctrl. Notice the ‘fingers’ of red KA dots extending from the central cluster of mixed Ctrl and KA data points. **(C)** Using a voxel gridding of the PC space (voxel size: 10 cubic units), we calculated the probability *P(KA)* for each event to be specific to KA treatment. The hourly IIS index of a mouse is the average number of detected events/h, weighted by their probability of being specific to the KA treatment. **(D)** Events with *P(KA) > 0.80*. Two visually identified groups (red ellipses) were separated along the PC1 axis (group a < −10 and group b > −10). **(E)** Each row shows example events from groups identified in D and from the remaining events in C. EEG amplitude was normalized and reported in terms of standard deviation (SD). **(F)** Average waveforms from group a (top), b (middle), and events with *P(KA)* < 0.80. Despite morphological differences, group a and b both show a wave following the detected spike, which is a known marker of epilepsy in TLE (Chauvière et al., 2012; Huneau et al., 2013).

Prior to EEG recordings, mice were subjected to an object-location novelty-recognition BPS task (**Methods – Mouse Behavior**) where their ability to discriminate between a moved object versus an identical unmoved object was measured for three different object shifts going from hard (7cm) to easy (21cm) (**Figure 3A**). Our results demonstrate that, on average, KA mice show lower mnemonic discrimination of object-location than control mice (**Figure 3B-C**). This impairment is most prominent for the object shift of intermediate difficulty (14 cm) which most control mice discriminated but most KA mice did not (**Figure 3B**). Control analyses (**Figure 3D-M**) revealed no side-preference bias (**Figure 3D**) and no significant impact of object exploration times (above a 4 s threshold) during sampling or testing on individual discrimination ability (**Figure 3E-F**). KA mice had no locomotor deficits (assessed through ambulatory activity, **Figure 3G-H**) and on average did not display higher anxiety levels (assessed by the time spent near walls) during the sampling and testing phases (**Figure 3K**). Ambulatory activity was slightly higher in KA mice during testing, and anxiety levels more variable, but there was no correlation between individual mnemonic discrimination ratios and ambulatory activity (**Figure 3I-J**) or anxiety levels (**Figure 3L-M**). These results suggest that mnemonic discrimination differences between KA and control mice in the BPS task were not confounded by individual motor ability, exploration or anxiety factors nor by differences of those factors between phases.

**Figure 3.**
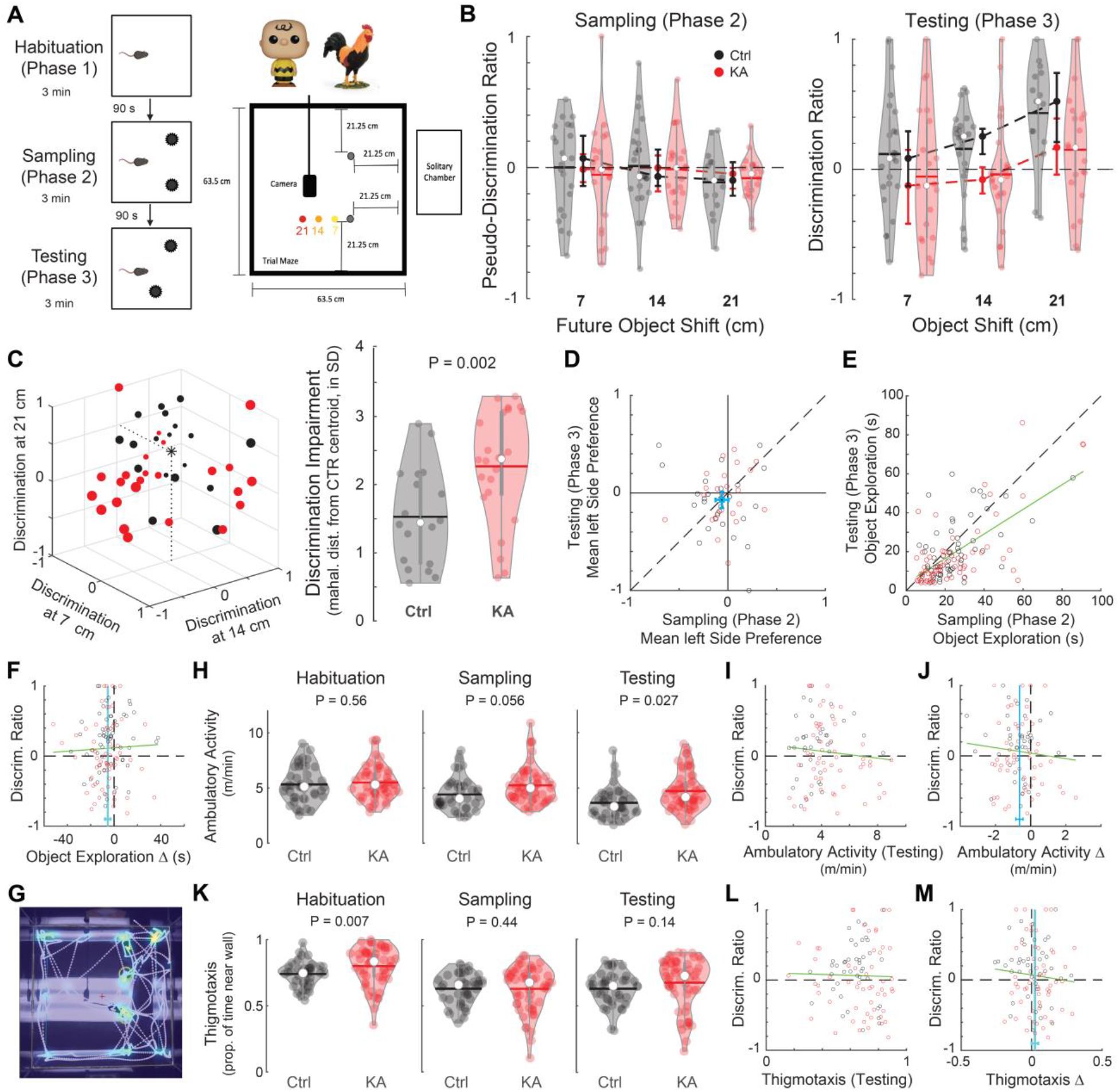
Kainate-injected mice have deficits in mnemonic discrimination. **(A)** Object location discrimination task (Left) and arena setup (Right). In Phase 1, a mouse was allowed to explore the empty square arena. In Phase 2, the same mouse was allowed to explore the same arena but with two new identical objects (Top right: example objects). In Phase 3, one randomly chosen object was shifted either 7, 14 or 21 cm from its original position and the mouse was allowed to explore again. Most mice performed the task once at each distance. Different objects were used for each distance and distance order was randomized, with a 1 week-interval. **(B)** Discrimination between the two identical objects, for each mouse and distance, in phase 2 (Left, sampling/control trial) and phase 3 (Right, actual discrimination test). Discrimination ratio = difference in exploration time for each object (moved - unmoved) normalized by the total object exploration time during a given phase. Mice with total object exploration times < 4 s during phase 2 or 3 are excluded. Violin plots show the distribution of discrimination ratios (open circle = median, horizontal bar = mean, scattered dots = trials). Solid circles with error bars between violin plots = bootstrapped 95% confidence interval of the median. In phase 2, mice do not discriminate between objects before they have moved (means not significantly different from 0). In phase 3, control mice (n = 27) prefer exploring the moved object when it has shifted enough (14 and 21 cm), whereas KA mice (n = 25) do not show significant discrimination on average. A two-way ANOVA with repeated measures confirms that mice discriminate better when objects are moved farther apart (Object Shift: *P* = 0.013, *F(1,123)* = 6.28), and that Ctrl mice discriminate better than KA mice (Treatment: *P* = 0.019, *F(1,123)* = 5.65). Interaction between Object Shift and Treatment was not significant and thus excluded from the ANOVA, but the bootstrapped confidence intervals of the medians suggest that difference between treatments is strongest at 14 cm. Note that phase 2 pseudo-discrimination ratios were not predictive of discrimination in phase 3 (linear regression: slope = 0.093, R^2^ = 0.4%, *P* = 0.5, *F(1,124)* = 0.49) and that a significant difference between treatments was still detected when considering the individual discrimination difference between phase 3 and phase 2 (Object Shift: *P* = 0.003, *F(1,123)* = 8.8453; Treatment: *P* = 0.0496, *F(1,123)* = 3.93). **(C)** An individual discrimination impairment score was computed for mice that were tested for all 3 object shift conditions (Ctrl: n = 18, KA: n = 24, no exclusion criteria). Left: Each animal is represented by a single dot with coordinates corresponding to its discrimination ratios in the 3 object shift conditions. The discrimination impairment was calculated as the Mahalanobis distance of each data point from the Ctrl centroid (black star) and is represented by the dot size: note that red dots (KA) appear larger than black dots (Ctrl). Right: Discrimination impairment scores are larger for KA than Ctrl animals (one-sided T-test: *P* = 0.002, *T(40)* = −3.04). **(D-M)** Control analyses. **(D)** Side-preference was computed for each animal as the exploration time difference between the left and right objects normalized by the total exploration time, averaged across Object Shift conditions. No side-preference was detected in either phases (cyan error bars: 95% CI of the means from two-sided one-sample T-tests. Phase 2: *P* = 0.07, *T*(51) = −1.87; Phase 3: *P* = 0.1 *T*(51) = - 1.69) nor treatment groups (ANOVA, Treatment: *P* = 0.39, *F*(1,50) = 0.75). **(E)** Total object exploration times in Phase 2 and 3 were correlated (green line: R^2^ = 42%, F(1, 124) = 88.4). There was no difference between object shift conditions or treatment groups (ANOVAs with repeated measures, Phase 2: Object Shift *P* = 0.39 *F*(1,123) = 0.74, Treatment: *P* = 0.83 *F*(1,123) = 0.05; Phase 3: Object Shift *P* = 0.65 *F*(1,123) = 0.20, Treatment *P* = 0.52 *F*(1,123) = 0.41). Sample size: 126 trials, as in Fig. B. **(F)** Difference in total exploration times (phase 3 – phase 2) was negative (mean: vertical cyan line, horizontal error bars: 95% CI from two-sided one-sample T-test, *P* < 0.0001, *T*(125) = −4.3), showing that animals explore objects 5.3 s less in phase 3 on average, likely due to habituation. Indeed, this total exploration time discrimination between phases was not predictive of Phase 3 mnemonic discrimination ratios (green line: R^2^ = 0.1%, *P* = 0.68, *F*(1, 124) = 0.17). There was no difference between object shift conditions or treatment groups (ANOVA with repeated measures: Object Shift *P* = 0.83 *F*(1,123) = 0.04, Treatment: *P* = 0.22 *F*(1,123) = 1.5). **(G-M)** Video records available from 43 mice (96 trials, same exclusion criteria as in B) were analyzed to extract animals’ trajectory and control for potential differences in ambulatory activity and anxiety levels. In H-M, individual mice are represented by 1-3 data points depending on how many shift conditions they were tested on and how many trials passed the exploration time criteria. Object Shift was never a significant effect. **(G)** video frame overlayed with detected trajectory in a sampling trial (color code for time spent at location: blue-to-yellow = low-to-high). Objects are highlighted by yellow circles. **(H)** Ambulatory activity (distance travelled / time spent traveling during trial) was used as a proxy for motor ability. KA mice showed no motor impairement and actually tended to move slightly more than Ctrl mice in phase 2 and 3 (ANOVAs with repeated measures, Phase 1: Object Shift *P* = 0.36 *F*(1,93) = 0.83, Treatment: *P* = 0.56 *F*(1,93) = 0.33; Phase 2: Object Shift *P* = 0.34 *F*(1,93) = 0.92, Treatment: *P* = 0.056 *F*(1,93) = 3.75; Phase 3: Object Shift *P* = 0.14 *F*(1,93) = 2.24, Treatment *P* = 0.027 *F*(1,93) = 5.06). **(I)** Ambulatory activity in phase 3 was not predictive of mnemonic discrimination ratios (green line: R^2^ = 0.8%, *P* = 0.40, *F*(1,94) = 0.72). **(J)** Ambulatory activity levels in phase 2 and 3 were well correlated (linear regression: R^2^ = 65%, P < 0.0001, F(1, 94) = 173) but activity in phase 3 was −63.7 cm/min lower (mean: vertical cyan line, horizontal error bars: 95% CI from two-sided one-sample T-test, *P* < 0.0001, *T*(95) = −6.1), likely due to habituation (consistent with F). Indeed, this discrimination between phases was not predictive of phase 3 mnemonic discrimination ratios (green line: R^2^ = 0.7%, *P* = 0.41, *F*(1, 94) = 0.68). There was no difference in ambulatory activity Δ between object shift conditions or treatment groups (ANOVA with repeated measures: Object Shift *P* = 0.73 *F*(1, 93) = 0.11, Treatment: *P* = 0.52 *F*(1,93) = 0.41). **(K)** Thigmotaxis was used as a proxy for anxiety levels. On average, KA and Ctrl mice had similar anxiety levels, with a slight but significant difference only in phase 1 (ANOVAs with repeated measures, Phase 1: Object Shift *P* = 0.79 *F*(1,93) = 0.07, Treatment: *P* = 0.007 *F*(1,93) = 7.6; Phase 2: Object Shift *P* = 0.98 *F*(1,93) = 0.0007, Treatment: *P* = 0.44 *F*(1,93) = 0.60; Phase 3: Object Shift *P* = 0.70 *F*(1,93) = 0.14, Treatment *P* = 0.14 *F*(1,93) = 2.17). The variance of anxiety levels was larger in KA animals, with clearly separate high and low anxiety groups visible in phase 3. Mice with the highest IIS indices (Fig. 2) were in the high anxiety group, KA mice with abnormally low thigmotaxis had lower IIS indices. We did not observe a clear relationship between thigmotaxis and ambulatory activity. **(L)** Thigmotaxis in phase 3 was not predictive of mnemonic discrimination ratios (green line: R^2^ = 0.4%, *P* = 0.53, *F*(1,94) = 0.39). **(M)** Thigmotaxis levels in phase 2 and 3 were well correlated (linear regression: R^2^ = 54%, P < 0.0001, F(1, 94) = 111) and barely different from each other (mean: vertical cyan line, horizontal error bars: 95% CI from two-sided one-sample T-test, *P* = 0.05, *T*(95) = 1.98). Thigmotaxis Δ (phase 3 – phase 2) was not different across shift conditions but slightly more pronounced in KA animals than Ctrl (ANOVA with repeated measures: Object Shift *P* = 0.86 *F*(1, 93) = 0.03, Treatment: *P* = 0.04 *F*(1,93) = 4.3). But thigmotaxis Δ was not predictive of mnemonic discrimination ratios (green line: R^2^ = 0.6%, *P* = 0.44, *F*(1, 94) = 0.60; KA only: R^2^ = 0.06%, *P* = 0.85, *F*(1, 94) = 0.03).

### DG computational impairments in TLE

To test our hypothesis that DG pathologies in TLE lead to a breakdown of the neural pattern separation function of the DG, which in turn would translate into the cognitive impairments we observed, we measured pattern separation in hippocampal slices from the same KA and control mice used above in behavioral and EEG experiments. Although the slice preparation isolates the DG from its natural inputs, it provides the advantage that applied inputs and recorded outputs can be known with certainty, a necessary condition for quantifying circuit-level pattern separation (Santoro, 2013; Madar et al. 2019a). Briefly, the assay has three steps (**Figure 4A-B** and see Madar et al. 2019a, b): 1) ensembles of stimulus patterns (simulating afferent input spiketrains) are generated, with known degrees of similarity to each other. These spiketrains are then fed into the DG by stimulating the lateral perforant path. 2) The response of a single GC is recorded in whole-cell current-clamp. 3) The similarity between the output spiketrains is compared to the similarity between the input spiketrains, revealing the degree of separation or convergence (**Figure 4C** and see **Material and Methods – Neural pattern separation analysis**).

**Figure 4.**
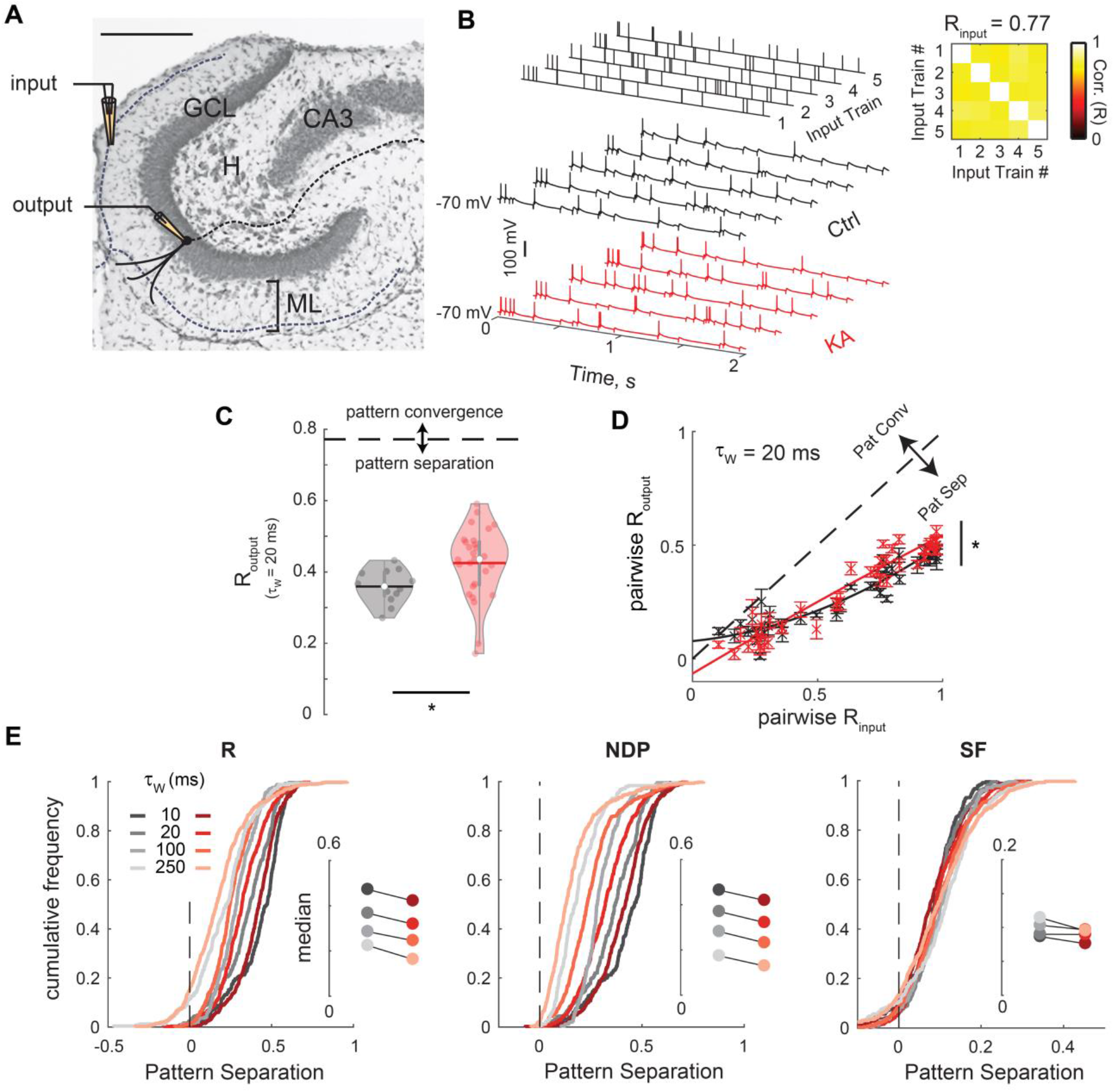
Epileptic mice have neuronal pattern separation deficits. **(A)** Histology of the DG in a horizontal slice (Cresyl violet/Nissl staining; scale bar: 250 μm), overlaid with a schematic of the experimental setup: a theta pipette in the ML is used to focally stimulate the outer molecular layer (input) while a responding GC is recorded via whole-cell patch-clamp (output). GCL: granule cell layer, H: hilus, ML: molecular layer. Solid lines represent dendrites and dashed lines axons. **(B)** Current-clamp recordings of the membrane potential of two different GCs (Ctrl and KA) in response to the same set of input trains. An input set is constituted of five different trains of electrical pulses following a Poisson distribution with an average rate of 10 Hz. The Pearson’s correlation coefficient (R) between two input trains is computed with a binning window (τw) of 20 ms (*Left*). R_input_ is the average of the ten pairwise coefficients, diagonal excluded. After converting the GC recordings to vectors of binned spike counts, the pairwise coefficients and their average (R_output_) can be computed the same way (Madar et al, 2019b). Note that the input set was repeated 10 times to yield output sets of fifty sweeps (only one repeat is shown). **(C)** Violin plots showing the distribution of GCs output correlation R_output_ (open circle = median, horizontal bar = mean). Data points correspond to single output sets (KA: 27 recordings from 24 GCs; Ctrl: 12 recordings from 11 GCs), all being responses to the same input set as in B.U-test comparing the medians: P = 0.011, Z = −2.541, rank sum = 156; two-sided two-sample KS test comparing the distribution shapes: P = 0.0037, D = 0.583. The dashed line corresponds to R_input_ (0.77): any value below the line implies effective pattern separation. Thus, GCs from kainate animals exhibit less pattern separation than controls. **(D)** Pattern separation was investigated over a wide range of input correlations by using four different input sets of five 10 Hz Poisson trains (at τ_w_ = 20 ms: R_input_ = 0.24, 0.45, 0.77, 0.95). Crosses and error bars correspond to mean +/- SEM (pw R_input_, pw R_output_) across multiple recordings (KA: 6-24 GCs per input set; Ctrl: 4-11 GCs). Data points below the identity line (dashed) correspond to pattern separation. The distributions between KA and Ctrl are significantly different (ANCOVA with separate parabolic models fitting the data points and not the means: P < 0.0001, F(3,714) = 15.485. Solid curves are the parabolic models used for the ANCOVA. **(E)** Levels of pattern separation measured using different similarity metrics (S: R, NDP and SF, see Methods) and timescales. Cumulative frequency distributions of the distance of (pw S_input_, pw S_output_) data points to the identity line in pattern separation graphs like in E. Positive values of the x-axis correspond to pattern separation, and negative values to pattern convergence. Insets show medians. For R and NDP, distributions are significantly shifted to the left, showing that GCs from KA exhibit less decorrelation and less orthogonalization (ANCOVA as in D, but using separate linear models: P < 0.01 for τ_w_ = 5 to 1000 ms. For R/NDP and τ_w_ = 10, 20, 100, 250 ms, P = <0.0001/<0.0001, <0.0001/<0.0001, 0.001/<0.0001, 0.011/0.0001, F(2, 716) = 15.2/18.2, 20.3/23.4, 6.87/15.97, 4.5/9.4). For SF, scaling levels are weakly but significantly shifted to less separation at large τ_w_ (SF: P > 0.2 except at 250 ms and 500 ms, with P = 0.026, 0.046, F(2, 716) = 3.65, 3.09 respectively).

We tested pattern separation levels in response to three types of input sets. Input sets of type 1 were constituted of Poisson spiketrains with a 10 Hz mean firing rate designed to have a prespecified average similarity as measured by the Pearson’s correlation coefficient (R) (**Figure 4B**). For an input set with spiketrains correlated by 77% between each other (timescale: τ_w_ = 20 ms), the output spiketrains of GCs from epileptic mice had a higher average correlation (R_output_) than GCs from control mice (**Figure 4C**). This demonstrates a decrease in pattern separation in the DG of epileptic mice. Exploring a wider range of input correlations shows that the pattern separation function of the DG of KA mice is generally impaired compared to the normal pattern separation function (**Figure 4D**). Furthermore, a decrease in pattern separation was observed at multiple timescales (millisecond to second) and using multiple similarity metrics that assume different neural codes (**Figure 4E**, see **Material and Methods – Neural pattern separation analysis**). The impairment is most noticeable using the NDP metric, showing that DG output patterns are less orthogonalized in KA animals at all timescales, whereas differences in pattern separation via scaling (SF metric) are small but significant at large timescales.

An input set of type 2, made of Poisson spiketrains similar in terms of R but with varying firing rates (**Figure 5A1**), and an input set of type 3, with 10.5 Hz uncorrelated trains (R = 0) with varying burstiness (**Figure 5B1**), were designed to explore a wide range of input similarities as measured by SF and to characterize DG computations on inputs with a variety of statistical structures (Madar et al., 2019b). As we have shown before, the normal DG exhibits low pattern separation via scaling for highly similar inputs, and significant pattern convergence for dissimilar inputs (Madar et al., 2019b). In KA mice, GCs show a decrease in both pattern convergence and separation via scaling: the DG computation function is steeper, closer to the identity line, meaning that the DG of KA mice is weaker at transforming the similarity of its inputs in general (**Figure 5A1 and B1**). When similarity is measured with R or NDP (binwise synchrony code) instead of SF (neural code focused on FR and bursting), these experiments confirmed that pattern separation is decreased in TLE (**Figure 5A2**) and that this deficit is strongest when inputs are highly similar (i.e., when pattern separation is theoretically most important) and weaker when inputs are already dissimilar (**Figure 5B2**).

**Figure 5.**
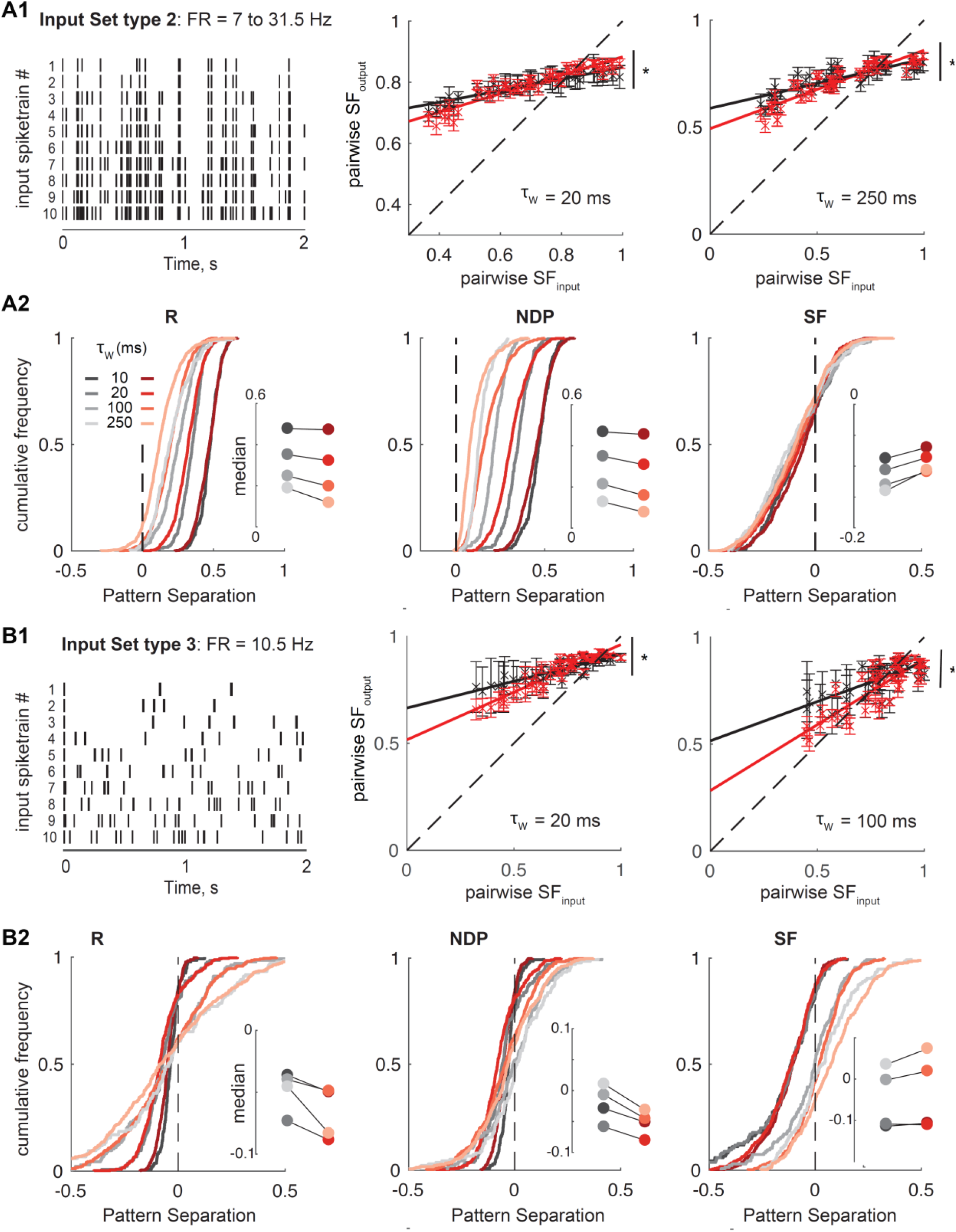
Multiple DG computations are affected by epilepsy. Our standard input sets (type 1, Figure 4) consisted of 10 Hz Poisson trains. Two other input sets (type 2 and 3) were designed to explore single GCs responses to inputs with diverse structures and statistics, both with a wide range of pairwise similarity as measured by SF (in contrast to input sets of type 1). **(A1)** *Left*: Input set type 2 was constituted of spiketrains following a Poisson distribution, each with a different firing rate (FR), but with an average R_input_ constrained around 0.75 (τ_w_ = 10 ms). *Middle and Right*: Pattern separation graphs showing the pairwise output spiketrain similarity as a function of the pairwise input similarity, as measured by SF with two different timescales, averaged across multiple GCs (KA: 16, Ctrl: 5). Distributions are significantly different, suggesting in particular that epilepsy causes a decrease in pattern convergence (via scaling) for low input similarities (ANCOVA with separate linear models, for τ_w_ = 20 and 250 ms respectively: P = 0.0016 and < 0.0001, F(2, 941) = 6.5 and 10.4). **(A2)** Levels of pattern separation for type 2 inputs, measured using different similarity metrics and timescales as in Figure 4. It confirms that epilepsy decreases the separation of similar Poisson input spiketrains as measured by R and NDP, consistent with Figure 4, and shows that differences in terms of SF, although small, are significant (ANCOVA as in A1: R: P < 0.0001 for τ_w_ = 20 ms up to 1000 ms, P = 0.02 and 0.18 for 5 and 10 ms; NDP and SF: P < 0.025 for τ_w_ up to 1000 ms. Detailed statistics for τ_w_ = 5, 10, 20, 50, 100, 250, 500 and 1000 ms respectively: R, P = 0.0243, 0.1820, <0.0001, <0.0001, <0.0001, <0.0001, <0.0001, <0.0001, F(2, 941) = 3.7, 1.7, 10.2, 15.1, 24.1, 36.0, 50.2, 27.0; NDP, P = 0.0022, 0.0240, <0.0001, <0.0001, <0.0001, <0.0001, <0.0001, <0.0001, F(2, 941) = 6.2, 3.7, 13.8, 17.4, 21.4, 26.0, 21.0, 16.8; SF, P = 0.0002, 0.0012, 0.0016, 0.0240, 0.0073, <0.0001, <0.0001, 0.0002, F(2, 941) = 8.7, 6.7, 6.5, 3.7, 4.9, 10.9, 10.4, 8.7). **(B1)** *Left*: Input set type 3 was constituted of spiketrains with 21 spikes (FR = 10.5Hz) that were distributed among bins to produce trains with varying burstiness (Madar et al., 2019b). R is close to 0 for all pairs. *Middle and Right*: same analysis as in A1 (KA: 8 GCs, Ctrl: 3 GCs). Distributions are significantly different, suggesting again that epilepsy causes a decrease in pattern convergence via scaling, for low input similarities (ANCOVA for τ_w_ = 20 and 100 ms: P < 0.0001 and F(2, 491) = 10.3 and 13.8 respectively). **(B2)** Same analysis as in A2. The directions of impairments are the same as in A2, showing that epilepsy decreases pattern separation in terms of R and NDP but slightly improves it in terms of SF. The shift via scaling is weak but significant at all timescales, and larger at the longest τ_w_. ANCOVA: R, P < 0.05 for τ_w_ = 5, 10 and 500 ms; NDP and SF, P < 0.007 for τ_w_ up to 500 ms. Detailed statistics for τ_w_ = 5, 10, 20, 50, 100, 250, 500 and 1000 ms respectively: R, P = 0.0451, 0.0004, 0.3727, 0.7357, 0.9340, 0.4271, 0.0012, 0.2889, F(2, 491) = 3.1, 8.1, 1.0, 0.3, 0.1, 0.8, 6.8, 1.2; NDP, P = <0.0001, <0.0001, 0.0003, 0.0012, <0.0001, 0.0069, <0.0001, 0.4154, F(2, 491) = 11.1, 24.3, 8.4, 6.8, 11.2, 5.0, 14.6, 0.9, SF, P = <0.0001, <0.0001, <0.0001, <0.0001, <0.0001, 0.0065, 0.0002, 0.0003, F(2, 941) = 42.1, 20.9, 10.3, 13.6, 13.8, 5.1, 8.5, 8.1.

### Pathological spiking patterns in a subset of GCs from epileptic mice

What features of GC output spiketrains are changed in TLE to explain the difference in DG computations between KA and control mice? On visual inspection, we noticed that many KA neurons occasionally fired short bursts of action potentials after a single input pulse (**Figure 6A**), which is unusual for control GCs (Madar et al., 2019a). To determine whether there was a difference in the spiking patterns of KA and control GCs, we measured three spiketrain features: 1) the average firing rate (FR) of a GC across a full recording set (fifty spiketrains), 2) the probability of spiking after a single input pulse (SP) and 3) the probability of bursting (p(Burst), i.e., more than one spike) after a single input pulse. Our results show that on average GCs from KA mice fire more faithfully after an input (**Figure 6B**) and also tend to fire in bursts (**Figure 6C-D**), which together lead to a higher FR (**Figure 6B, E**).

**Figure 6.**
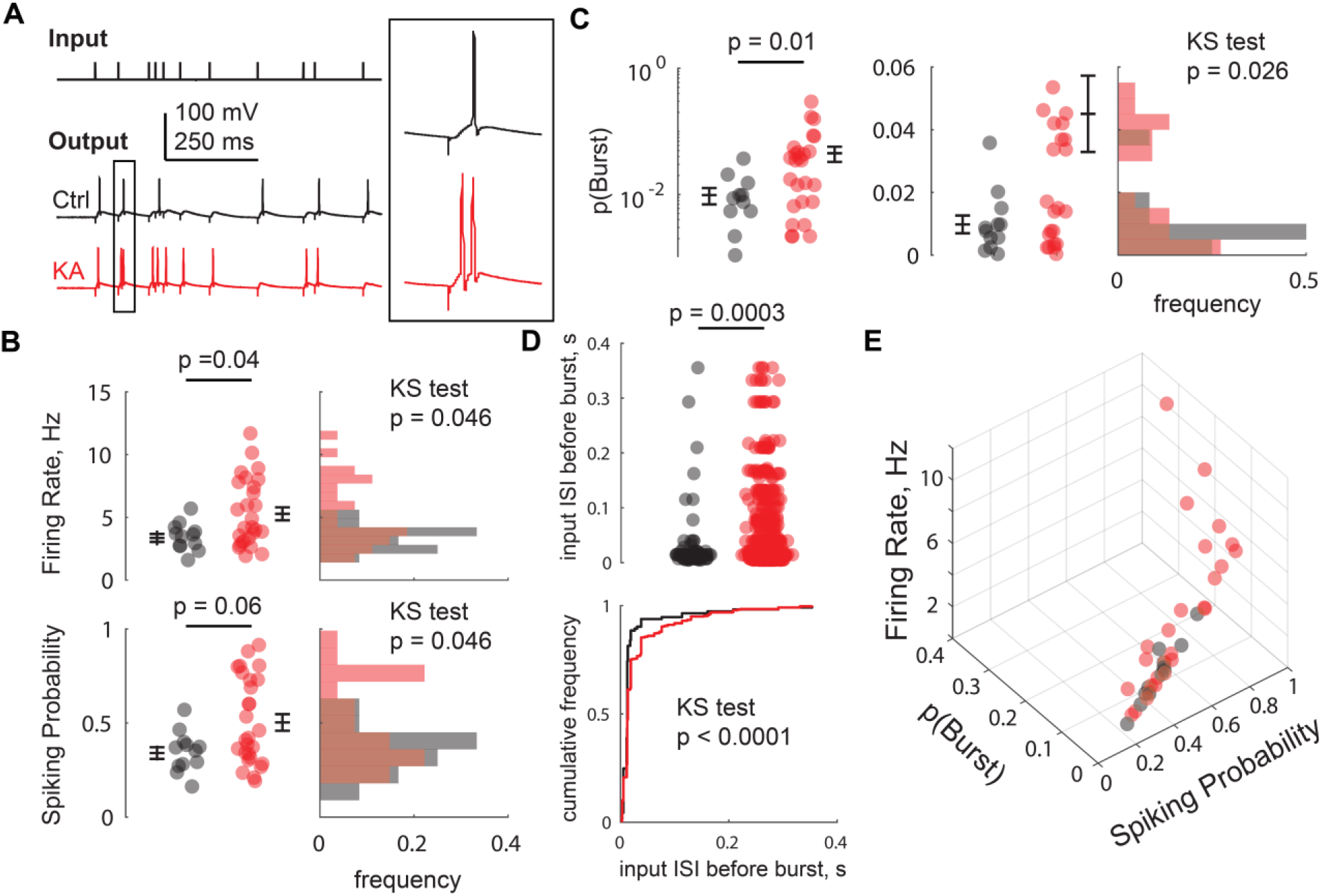
In epileptic mice, a subpopulation of GCs shows pathological spiking patterns. **(A)** Example of current-clamp recordings in GCs from a kainate-injected (KA) vs a control animal in response to the same Poisson input train, illustrating that some GCs from KA spiked more, and had a tendency to fire short bursts (2-4 spikes riding a single EPSP, see inset). **(B-E)** Some GCs from KA exhibited larger firing rates due to a higher probability of spiking at least once after an input spike (spiking probability), and sometimes a higher probability of spiking more than once between two input spikes (p(Burst)) than GCs from controls. The firing rate, spiking probability and p(Burst) of a neuron were all computed as the average over the fifty sweeps of an output set from a pattern separation experiment (input set type 1, see Figure 4B). **(B)** *Left*: data points correspond to the same recordings as in Figure 4C (KA: 27; Ctrl: 12). Black dash and error bars are mean +/- SEM. A U-test was used to compare the medians, showing that on average FR and SP are higher in KA (FR: P = 0.0415, Z = −2.04, rank sum = 172.5; SP: P = 0.0613, Z = −1.87, rank sum = 168.5). *Right*: Frequency distributions of the same data. A one-sided two-sample KS test shows that the KA distribution has a larger tail in both cases (FR and SP: P = 0.0465, D = 0.4074), indicating that a subset of KA GCs are pathological. **(C)** Same as in B for p(Burst). Note that the left graph has a log10 scale showing all data points, whereas the middle graph has a linear scale zoomed in (i.e. not showing the 5 largest KA values). The graph on the right has the same scale as the middle graph. Together, they suggest that in KA, there might be a healthy population of GCs coexisting with a different population of pathologically bursty GCs (U-test: P = 0.0153, Z = −2.16, rank sum = 168.5; KS test: P = 0.0259, D = 0.4444). **(D)** Some bursts were detected by our algorithm in GCs from Ctrl mice, but the vast majority of those were due to the temporal summation of two EPSPs resulting from input spikes occurring close in time. In contrast, a large number of bursts in GCs from KA mice come after an isolated input spike. *Top*: Time interval between the two input spikes (inter-spike-interval, ISI) preceding a given burst. Data points correspond to all detected bursts from the recordings in C. U-test: P = 0.0003, Z = −3.58, rank sum = 5906.2. *Bottom*: Cumulative frequency distributions of the same data. A one-sided two-sample KS test demonstrates that the distributions are different (P < 0.0001, D = 0.2699). **(E)** Same data as in B and C: the elbow in the distribution of SP, p(Burst) and FR values visually defines two clusters of neurons. One cluster contains GCs from both KA and Ctrl mice (with GCs from both groups spanning a large range of SP values but low p(Burst)) and can be considered “normal”. In contrast, the other cluster, with highest SP combined with high p(Burst), contains only GCs from KA animals and can thus be considered “pathological”.

Closer inspection suggests that the distributions of SP, p(Burst) and FR are different between KA and controls, with larger variance and upper tails for GCs from KA mice **(Figure 6B-E).** Although we did not formally test for discrete clusters, visual inspection of Figure 6E shows that some GCs from KA animals have higher SP, p(Burst) and FR than any control GC. Thus, it appears that, following an epileptogenic insult, only a subset of GCs displayed pathological characteristics amidst a background of seemingly normal GCs.

What is the origin of the pathological spiking patterns? We first controlled that stimulation intensities were comparable between treatment groups and were not causing the abnormal spiking patterns (**Figure 7A**). We next asked whether the abnormal firing came from changes in the synaptic drive. To answer this question, we measured the synaptic input-output relation (Ewell and Jones, 2010) by recording GCs first under current-clamp then under voltage-clamp, in response to the same pattern separation stimulus protocol (**Figure 7B1-2**). Our data show that the average excitatory drive per neuron was not different between KA and control groups, and that it does not predict burstiness (**Figure 7B3**). A finer grained analysis on the individual currents suggests that although the excitatory drive and the spiking output are correlated, high amplitude EPSCs are neither sufficient nor necessary to elicit a burst (**Figure 7B4**). Bursting does not seem to result from other changes in excitatory input-output coupling either (**Figure 7B5**), which suggest that inhibition might be involved. GCs are indeed subjected to strong tonic, feedforward and feedback inhibition that controls the sparseness of their activity (Coulter and Carlson, 2007; Ewell and Jones, 2010; Pardi et al., 2015; Lee et al., 2016) and is altered in TLE (Alexander et al., 2016; Dengler and Coulter, 2016). We did not record evoked IPSCs directly but, in normal mice, partial block of inhibition elevates GC firing rates and causes bursts (Madar et al., 2019b) similar to what we observed here in KA mice. Although indirect, our results thus converge to support the hypothesis that pathological spiking arises from an excitation/inhibition imbalance in a subset of GCs, likely driven by a decrease in inhibition.

**Figure 7.**
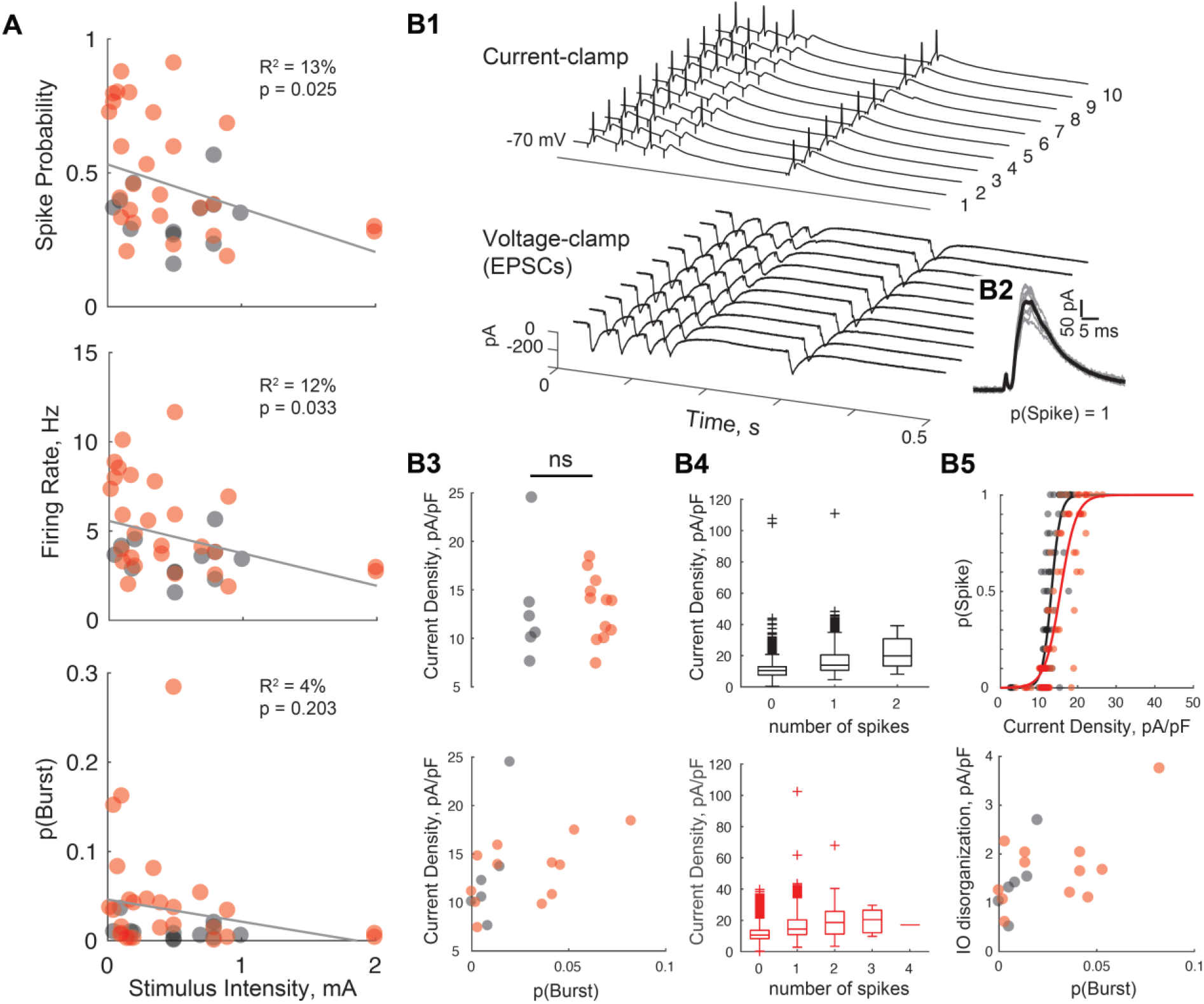
Pathological bursting is not explained by aberrant excitation. **(A)** Bursting and high firing rates were not an artefact of stimulation differences: The current intensity of the electric stimulation did not significantly differ between KA and Ctrl (same recordings as in Figure 4C and 6B-E, U-test: P = 0.2991, Z = 1.04, rank sum = 274.5) and stimulus intensity was a poor predictor of pathological firing patterns (as indicated by the low R^2^ values of linear regressions. For FR/SP/pBurst: R^2^ = 12.75%/11.69%/4.35%, F(2,37) = 5.4/4.9/1.7, P = 0.0257/0.0332/0.2027). **(B)** For a subset of neurons in A (KA: 12; Ctrl: 6), we followed the current-clamp recording with a voltage-clamp recording (Vhold = −70mV) in response to the same pattern separation protocol in order to assess the excitatory synaptic drive and its relationship with spiking behavior. The stimulus electrode location and current intensity was unchanged. **(B1)** Example of current-clamp (top) and voltage-clamp (bottom) recording in the same GC from a Ctrl animal. Only the first 500 ms of the ten responses to repetitions of the first input train shown in Figure 4B are displayed. *Top*: action potentials are truncated at 0 mV. *Bottom*: stimulation artefacts and occasional unclamped spikes were blanked. **(B2)** The ten EPSCs (grey) associated to first pulse of input trains in B1 and the mean EPSC (black). We assessed the excitatory drive as the maximum inward current in the interval between each input pulse minus the current baseline (i.e., mode of the current over the full sweep). P(Spike) was defined as the probability of spiking during this interval across the ten sweeps of the current-clamp recording. **(B3)** *Top*: The mean EPSC density for each neuron was not significantly different between KA and Ctrl (U-test: P = 0.5532, rank sum = 50; two-sided two-sample KS test: P = 0.1935, D = 0.4167), suggesting that pathological spiking cannot be explained by larger EPSCs in KA mice. *Bottom*: Indeed, no clear relationship exists between p(Burst) and the mean current density, as some GCs with a great synaptic drive do not exhibit much bursting, and vice versa. **(B4)** Consistently, the peak current density associated to each inter-input-interval in a recording is not well correlated to the corresponding number of spikes in the current-clamp trace, for both Ctrl (top) and KA (bottom). For example, excitatory currents of the same magnitude can be associated to 0, 1, 2, 3 or 4 spikes. EPSC amplitude is thus not sufficient to explain the spiking output, which suggests that other factors, like reduced inhibition, are likely implicated in causing pathological bursts. **(B5)** *Top*: The average EPSC density between two input spikes is plotted against the corresponding probability of spiking p(Spike), as in Ewell and Jones (2010). The resulting input-output (IO) distribution is fitted with a sigmoid (two GCs representative of KA and Ctrl are shown). *Bottom*: In a subset of GCs, mostly from KA, The IO distribution appears disorganized and sometimes difficult to fit. A proxy of this disorganization is the standard deviation (SD) of the current density, averaged across each level of p(Spike) excluding p(Spike) = 0 and 1. The average SD for a given GC was plotted against the propensity of that neuron to fire bursts. There is no clear relationship between these two quantities, suggesting that: 1) decoupling between the excitatory drive and spiking probability is not directly related to bursting and 2) a subset of GCs from KA with apparently healthy spiking patterns could already exhibit a more subtle but pathological IO disorganization.

How do high FR and bursts relate to the DG computational impairment measured in KA mice? We first tested the hypothesis that higher GC firing rates and burstiness yield less pattern separation as measured with R (**Figure 8A**). In GCs of young mice, FR and pattern separation are related, but loosely (Madar et al., 2019a), whereas here in adult mice there is a strong linear relationship, such that GCs with pathologically high FR exhibit pathologically low pattern separation. Similarly, abnormal bursting corresponds to abnormally low levels of pattern separation. This analysis relates cell-wise spiking features (FR or pBurst averaged across all spike trains of a recorded neuron) to a form of pattern separation that is not theoretically concerned with such features (Madar et al., 2019b).

**Figure 8.**
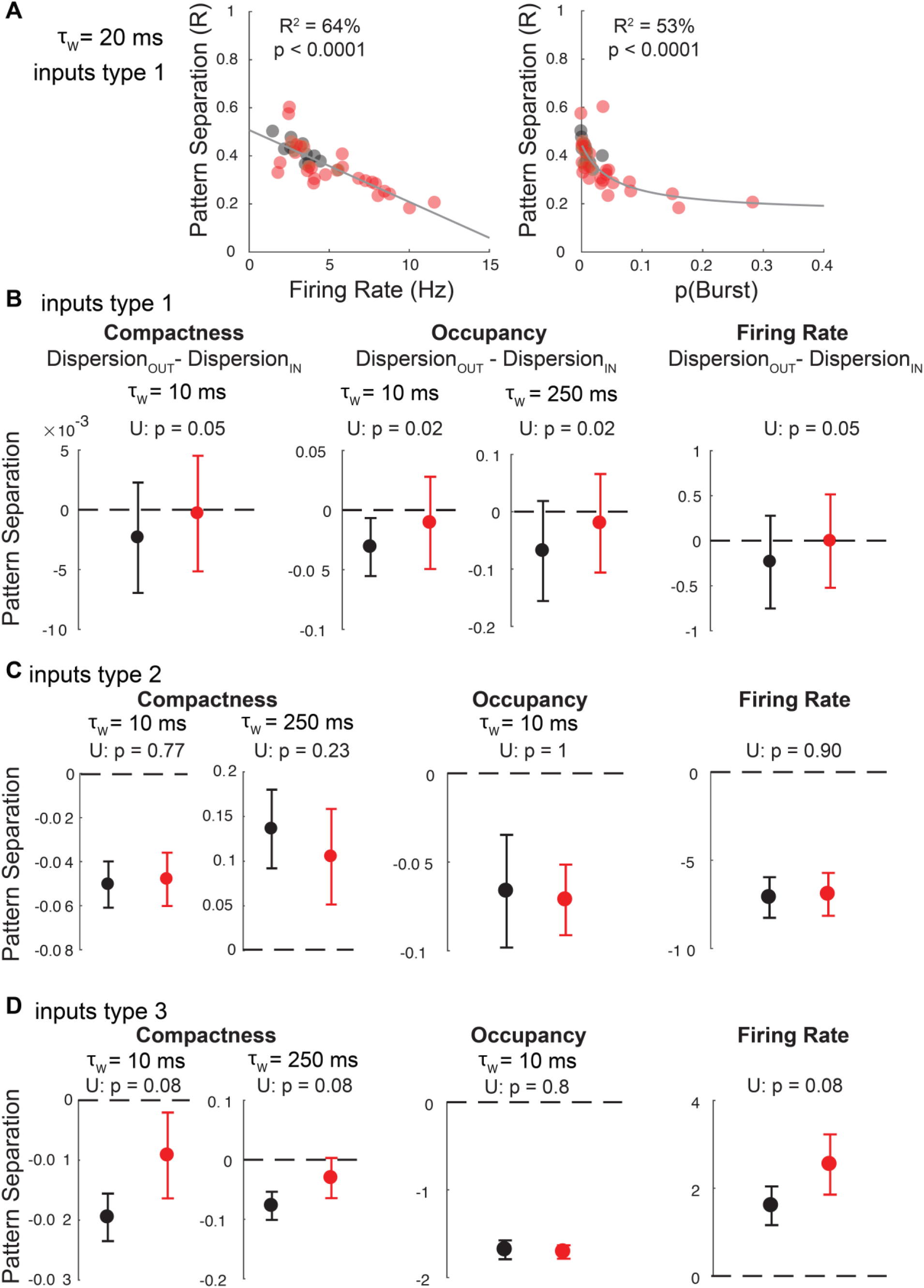
Pathological spiking patterns explain pattern separation differences in epilepsy. **(A)** Neurons with high average firing rates and burstiness exhibit lower pattern separation (computed as R_input_ – R_output_). Same data as in Figure 4C and 6B-C. There is a strong linear relationship between firing rates and decorrelation (Grey line, R^2^ = 63.68%, F(2,37) = 64.9, P < 0.0001). The relationship between p(Burst) and decorrelation is better described by an inverse function with a horizontal asymptote (see Methods – Software and statistics): R^2^ = 53.1%, R^2^ adjusted for 3 parameters = 50.5%, F(3,36) = 20.4, P < 0.0001. **(B-D)** Mean +/- SEM pattern separation levels across recordings when considering burstiness codes (Compactness or Occupancy) or a rate code (spiking frequency across a 2s sweep) instead of the binwise synchrony code assumed by R. Compactness is the proportion of time bins with at least one spike, whereas occupancy is the average number of spikes in a bin. Pattern separation corresponds to more dispersion in compactness, occupancy or firing rate in the output spiketrains than the input spiketrains. Negative values mean there is pattern convergence. U-tests comparing medians of GCs from KA and Ctrl groups were performed (p-values in panel). **(B)** In response to input sets of type 1 (10 Hz Poisson trains, same recordings as in A and Figure 4), GCs exhibit low levels of convergence, if any, in terms of compactness, occupancy and rate codes, and these levels are slightly but significantly shifted to less convergence in GCs from KA for τw = 10 ms. This means that the relationships between firing rate or burstiness and pattern decorrelation observed in A is not due to variations of the firing rate or burstiness across sweeps. Detailed statistics at 10, 20, 50, 100, 250 ms, Compactness: P = 0.0482, .1892, .1628, .2925, .7584, Z = −1.97, −1.97, −1.31, −1.39, 0.31, rank sum = 745, 745, 801, 794, 939; Occupancy: P = 0.0177, 0.2586, 0.3624, 0.1425, 0.0211, Z = −2.37, −1.13, −0.91, −1.47, −2.31, rank sum = 712.5, 816.5, 835, 788, 717; FR: P = 0.0553, Z = −1.92, rank sum = 750. **(C)** In response to an input set of type 2 (Poisson trains with different firing rates, same recordings as in Figure 5A), GCs from KA and Ctrl do not show significant differences (P > 0.2 for all timescales and codes). Detailed statistics at 10, 20, 50, 100, 250 ms, Compactness: P = 0.7726, 0.7726, 0.8365, 0.9014, 0.2312, Z = −0.29, −0.29, −0.21, 0.12, 1.20, rank sum = 51, 51, 52, 57, 70; Occupancy: P = 1, 1, 0.9671, 0.5915, 0.2006, Z = 0, 0, 0.04, −0.54, −1.28, rank sum = 55, 55, 56, 48, 39; FR: P = 0.9014, Z = −0.12, rank sum = 53. **(D)** In response to an input set of type 3 (10 Hz trains with varying compactness and occupancy, same recordings as in Figure 5B), GCs exhibit convergence in terms of compactness and occupancy and separation in terms of rate codes. Only computations in terms of compactness and firing rate are mildly shifted in GCs from KA for τw = 10 ms. Detailed statistics at 10, 20, 50, 100, 250 ms, Compactness: P = 0.0848, rank sum = 9; Occupancy: P = 0.8, 0.7758, 0.9212, 0.3758, 1, rank sum = 19.5, 20, 17, 23, 18; FR: P = 0.0848, rank sum = 9.

Other forms of pattern separation, more directly related to FR or bursting, can in theory also be performed: for example by increasing spiketrain-to-spiketrain *variability* in FR or burstiness of the output neuron, even if the average quantities were identical (Madar et al., 2019b). Because pathological GCs only occasionally fired bursts, we asked whether TLE could affect such forms of pattern separation. To test this, we measured the spiketrain-to-spiketrain variability in FR as well as in two complementary measures of burstiness (Compactness and Occupancy, see **Materials and Methods – Neural pattern separation analysis**). For input sets of type 1 (10 Hz Poisson), GC output variability was slightly lower in terms of burstiness or FR. This weak type of pattern convergence disappeared in GCs from KA mice (**Figure 8B**). For input sets of type 2 and 3, no significant difference between KA and Ctrl was detected (**Figure 8C-D**). Overall, Figure 8 suggests that TLE reduces DG neural pattern separation mostly by raising average levels of FR and burstiness, rather than through spiketrain-to-spiketrain variations of those features.

### Electrographic, behavioral and computational pathologies in individual mice

We have shown that, on average, KA mice develop EEG abnormalities (**Figure 2**), suffer mnemonic discrimination impairments (**Figure 3**) and have a bursting subpopulation of GCs with pattern separation deficits (**Figures 4–8**). Because we often performed all of the aforementioned experiments in the same mice, we next asked how those different epilepsy-related pathologies are linked at the individual level (**Figure 9**). In a simple framework where behavioral impairments are caused by computational deficits that are due to reorganization of DG network function that also leads to an increase in epileptiform EEG events, one could expect a simple relationship between all the variables we measured. Interestingly, our data suggest a more complex view. For example, animals with few interictal spikes can harbor some pathological neurons (**Figure 9A**) and be highly impaired at the BPS task (**Figure 9B**). This shows that both DG network pathologies and mnemonic impairments can occur independent of, or before, early EEG abnormalities. Conversely, both normal and pathological GCs were recorded in mice with more advanced EEG pathology, which confirms that those two subpopulations can coexist in the same KA animal (**Figure 9A**). Mice with clearly epileptiform EEG activity can also sometimes perform in the normal range on the BPS task (**Figure 9B**). As expected from theory we found a correlation between neuronal pattern separation and mnemonic discrimination (**Figure 9C**), but only for the behavioral test with medium interference levels, which had led to the most salient discrimination difference between treatments (**Figure 3B**) and is most likely to require DG’s integrity and correct functioning (Engin et al., 2015; Pofahl et al., 2021). When considering behavioral performance over all interference levels, the variability in individual behavior and in cell sampling prevents us to definitively conclude, but we note that KA animals with the largest cognitive impairments all had at least one recorded GC with pathological firing and decreased pattern separation (**Figure 9C**). Finally, in cases where EEG, BPS and patch-clamp data were all obtained from each animal, the combination of these measurements yielded a very obvious separation between normal and KA subjects (**Figure 9D**). Note that considering an EEG index based on individual clusters of IISs identified in Figure 2D did not change any of the conclusions stated above. Overall, our results suggest that epilepsy-related pathologies do not all develop in concert and that computational, behavioral and electrographic measures provide complementary informative dimensions that, together, better assess the epileptic state than any single dimension or pair of dimensions.

**Figure 9.**
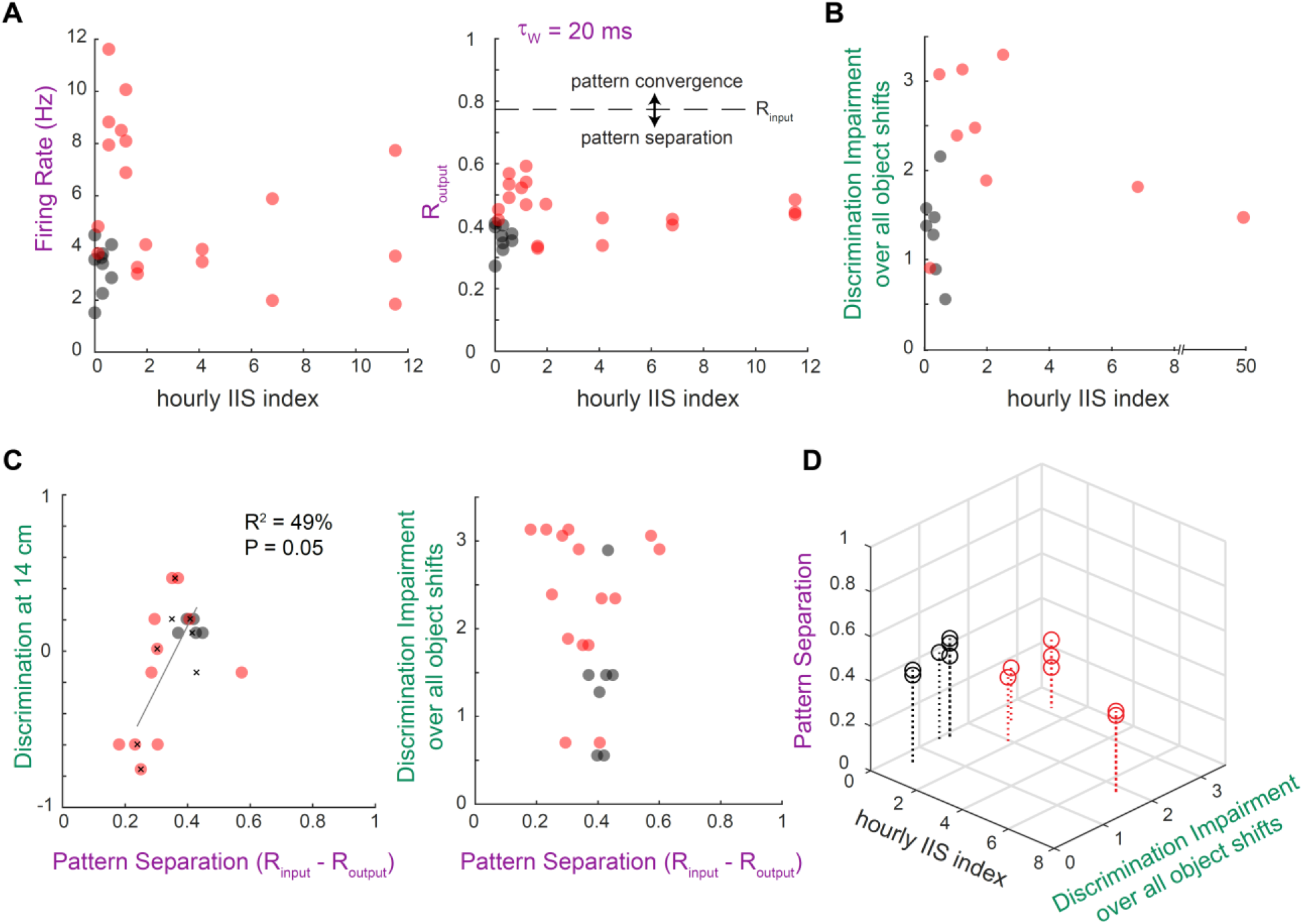
EEG, behavioral and computational measures provide complementary insights about epileptic pathology in individual animals. **(A-D)** Scatter plots relating the electrographic measure of IISs developed in Figure 2 (black axis labels), measures from patch-clamp recordings of individual GCs from the same animals (purple axis labels) and behavioral discrimination measures described in Figure 3 (green axis labels). Not all three types of records are available for every animal, which leads to differences in the samples shown in each panel. **(A)** Data points correspond to individual GCs from animals in which both EEG and patch-clamp data were recorded. Each animal has a unique average hourly IIS index value. Only a subset of GCs are outside of the Ctrl range (i.e. pathological). For example, in the mouse with the highest IIS index (~12), only one of the three recorded GCs exhibited a pathological FR (~8 Hz, left panel) and all had a similar R_output_ slightly above the Ctrl range (~0.5, right panel). In contrast, some GCs with abnormally high FR (~6-12 Hz)_and R_output_ (~0.5-0.6) came from animals with low IIS index (~0-2). **(B)** Data points correspond to individual animals in which both EEG and behavior were measured. Discrimination Impairment is the Mahalanobis distance from Ctrl centroid computed in Figure 3C. In KA animals, low IIS index can correspond to either normal or abnormal mnemonic discrimination, and despite having a high IIS index some animals have normal discrimination. Results were qualitatively similar using the discrimination ratios computed in Figure 3B. **(C)** Data points correspond to individual GCs from animals in which both behavior and patch-clamp data were recorded. *Left*: Black crosses represent the average neuronal pattern separation for each animal. These averages were used for linear regression: mnemonic discrimination ratios after a 14 cm object shift (Fig. 3B) and DG pattern separation (Fig 4C) were correlated (grey line: R^2^ = 49%, *P* = 0.05, *F*(1,6) = 5.75). No obvious relationship was observed for the other conditions (7 and 21 cm) but all KA mice had at least one pathological GC. *Right*: Same analysis for mice tested on all 3 object shift conditions (data from Fig 3C). The relationship is unclear, but all KA mice with large impairments had at least one pathological GC. **(D)** Data points correspond to individual GCs from animals in which EEG, behavior and patch-clamp data were all collected. Stems show which GCs were recorded from each animal. The lower right corner would correspond to the highest pathology in electrographic, behavioral and computational categories. Despite any heterogeneity or apparent ambiguity when only one or two of these measures are considered (above), when all three measures are available for each animal, the KA group is very clearly separable from the Ctrl group.

## Discussion

### Cognitive and computational deficits in TLE

We provide the first experimental evidence that TLE is characterized by both mnemonic discrimination impairments and neuronal pattern separation deficits.

Our data in human patients (**Figure 1**) are consistent with recent studies showing that TLE leads to mnemonic discrimination impairments (Reyes et al., 2018; Poch et al., 2020). Reyes and colleagues employed a spatial memory task whereas Poch and colleagues used an object-recognition task akin to ours. However, the specific task that we used is known to be DG-dependent in humans (Baker et al., 2016; Stark et al., 2019) and the TLE-related deficits observed in this task (see also Lalani et al., 2021) were not explained by DG-independent recognition impairments (**Figure 1E**), allowing us to infer a potential role of DG pathologies and further investigate these in mice.

Our behavioral experiment in mice (**Figure 3**) also complements previous studies (Bui et al., 2018; Kahn et al., 2019) by testing short-term (minutes) rather than long-term (24h) object-location memory and by testing multiple, parameterized levels of mnemonic interference, as recommended for rigorous testing of mnemonic discrimination (Hunsaker and Kesner, 2013; Liu et al., 2016). Although not demonstrated, the DG-dependency of our task in mice is likely because: 1) variants of this task are DG-dependent (Bui et al., 2018; Kahn et al., 2019; Pofahl et al., 2021), 2) DG supports discrimination for short to long term memory, as long as the task is not too easy or difficult (Gilbert et al., 2001; Engin et al., 2015; Pofahl et al., 2021), 3) we tested several degrees of difficulty and found an overall deficit in TLE that was most salient for the intermediate level of difficulty. Our results and the literature together suggest that mnemonic discrimination deficits are shared by humans and mice with TLE, occur across different memory domains and modalities, and point to DG malfunctions.

The impact of TLE-related DG pathologies on neural pattern separation had so far been investigated only in a computational model (Yim et al., 2015). This model suggested that mossy fiber sprouting and increased synaptic transmission from the perforant path to GCs could lead to a breakdown of pattern separation. We experimentally confirmed that TLE leads to a deficit in neural pattern separation (**Figure 4**). However, the deficit we observed is not as large, possibly because the model 1) considered synaptic transmission as deterministic rather than probabilistic and plastic, 2) assessed a different form of pattern separation (patterns were defined as population ensembles of very short spiketrains, thus focusing on a population code whereas we investigated temporal codes) and 3) considered severe degrees of epilepsy.

### The role of sparsity in neuronal pattern separation and DG gating

We found that GCs from epileptic mice exhibit pathological spiking consistent with a breach of the DG gatekeeper function (**Figure 6**) that leads to decreased neuronal pattern separation (**Figure 8A**). As noted by others, the gating and pattern separation functions of DG may be related (Dengler and Coulter, 2016) because the filtering of incoming cortical activity into a sparser output can be a mechanism underlying pattern separation based on population codes (O’Reilly and McClelland, 1994; Dieni et al., 2016; Cayco-Gajic and Silver, 2019; Braganza et al., 2020). Our results suggest that maintaining a sparse output also supports pattern separation of spiketrains in the time domain, at the level of single GCs.

Rare investigations of the spiking output of single GCs in epileptic tissue have reported that single stimulation of the perforant path sometimes leads GCs to fire bursts of spikes (Lynch et al., 2000; Kobayashi and Buckmaster, 2003). We expand on this by showing that bursts also emerge in response to complex, naturalistic stimulation patterns (**Figures 6 and 7**). It is important to note that occasional bursting is observed in normal GCs in vivo (Pernía-Andrade and Jonas, 2014), but this is very unusual in slices (Mongiat et al., 2009; Ewell and Jones, 2010; Zhang et al., 2012; Dieni et al., 2016; Madar et al., 2019a). In our experiments, bursting GCs therefore demonstrate a breach of the dentate gate. Because bursts are known to be more efficient at driving spiking in CA3 pyramidal neurons (Henze et al., 2002), a single GC firing bursts will have a higher probability of making downstream CA3 pyramidal cells fire. This could have deleterious effects on memory encoding and promote seizures, as more active CA3 neurons would 1) increase the chance of overlap between memory representations (Madar et al., 2019a, 2019b; Lee and Lee, 2020) and 2) overexcite a recurrent excitatory circuit, a mechanism of seizure generation (Traub and Wong, 1982; Santhakumar et al., 2005; de la Prida et al., 2006). Thus, our findings are consistent with the idea that pathological firing or bursting of even a subset of GCs could simultaneously contribute to deficits in mnemonic discrimination and epileptiform activity in TLE.

Our study also provides new insight on the mechanisms underlying GCs increased ability to burst by testing the influence of the excitatory drive (**Figure 7B**). The main source of excitation to GCs, the perforant path, is sometimes assumed to be augmented in TLE (Yim et al., 2015) given evidence of increased release probability at the lateral perforant path synapse (Scimemi et al., 2006). Our results do not directly contradict this, but we did not find a difference in the average synaptic current evoked in GCs by Poisson input spiketrains applied to the perforant path (**Figure 7B3**). More work is required to carefully test potential TLE-related changes in the coupling between perforant path and GCs, as well as EPSC kinetics and short-term dynamics (Madar et al., 2019a), but it is clear that the size of excitatory drive alone is not a good predictor of bursting propensity in GCs (**Figure 7B3-4**). We also previously showed that partial blockade of inhibition can produce GC bursting and pattern separation deficits (Madar et al., 2019b). Thus, in accordance with past research (Kobayashi and Buckmaster, 2003), we suggest that the strong decrease in synaptic inhibition characteristic of TLE (Dengler and Coulter, 2016; Dengler et al., 2017) is a likely driver of pathological bursting.

Perhaps the most intriguing insight we bring on the failure of the DG as a gate and pattern separator in TLE is that this failure occurs only in a subset of GCs, upon a background of apparently normal GCs in the same epileptic network. This finding invites many more questions: In the epileptic DG, is there a spectrum of spiking patterns among GCs going from normal to pathological, or are there two discrete types? What is the ratio of normal to pathological GCs and does it evolve during epileptogenesis? Do GCs become pathological due to changes in intrinsic properties or networking? Our results suggest deficits in inhibition, but additional mechanisms could be in play (Kelly and Beck, 2017). Anatomical or biophysical differences between GCs could be related to different birthdates of adult-born GCs relative to an epileptogenic brain insult, resulting in only a subset of GCs integrating abnormally into the DG network (Kron et al., 2010). Alternatively, pathological GCs could develop from a more susceptible and active subclass of mature GCs (Erwin et al., 2020). In any case, the discovery that the GC population in the epileptic DG is not homogeneous in terms of spiking patterns is a crucial step towards developing new treatments that would specifically target pathological neurons to avoid deleterious side-effects.

### The link between mnemonic discrimination, pattern separation and TLE

For thirty years the DG has been hypothesized to support mnemonic discrimination by performing neuronal pattern separation (Treves et al., 2008), but despite important efforts (McHugh et al., 2007; Marrone et al., 2011; Ramirez et al., 2013; Yokoyama and Matsuo, 2016; Allegra et al., 2020; Woods et al., 2020), no experiment has yet directly connected DG input-output computations with their role in memory. Here, we confirmed our previous work showing that the DG itself performs temporal pattern separation (Madar et al., 2019a, b) and we showed that a deficit in pattern separation is associated with a deficit in mnemonic discrimination, on average (Figure 3, 4) and at the individual level (Figure 9C). This is correlative evidence, linking non-simultaneous experiments, we thus cannot exclude that other DG or non-DG processes (Bernier et al., 2017; Guise and Shapiro, 2017; Hainmueller and Bartos, 2020) are impaired in KA mice and partially lead to the observed cognitive deficits. A full demonstration of the causal link between DG pattern separation and mnemonic discrimination will require measuring and manipulating DG computations in vivo during episodic memory tasks, but our study is a preliminary step toward understanding how DG circuit-level computations relate to a high-level cognitive function and how these processes fail in TLE. A mechanistic understanding of this relation may lead to new early diagnosis tools (e.g. cognitive tests like in Figure 1) as well as therapies alleviating memory disorders in TLE. For example, our finding that a subset of GCs with pattern separation deficits can develop without EEG hallmarks of epilepsy (Figure 9A) suggests a mechanism for memory impairments that often occur early during epileptogenesis (Jones et al., 2007; Chauvière et al., 2009; Witt and Helmstaedter, 2015). Similarly, the coexistence of pathological and normal neurons could explain memory impairments in patients without hippocampal sclerosis (Rayner et al., 2019).

## Acknowledgements

We thank Lin Lin who helped start behavioral tests in mice in the Jones lab and Drs. C.E.L Stark and M.A. Yassa for allowing us to use software and materials they developed. This work was supported by the US National Institutes of Health (MVJ: RO1 NS075366), the University of Wisconsin Institute for Clinical and Translational Research (MVJ.; NIH/NCATS UL1TR000427), the US Department of Defense (RKM: PR161864) and Lily’s Fund for Epilepsy Research (ADM.; 2015 fellow).

